# Myosin IIA interacts with the spectrin-actin membrane skeleton to control red blood cell membrane curvature and deformability

**DOI:** 10.1101/202556

**Authors:** Alyson S. Smith, Roberta B. Nowak, Sitong Zhou, Michael Giannetto, David S. Gokhin, Julien Papoin, Ionita C. Ghiran, Lionel Blanc, Jiandi Wan, Velia M. Fowler

**Author notes:** These authors contributed equally (Alyson S. Smith, Roberta B. Nowak).

## Abstract

The biconcave disc shape and deformability of mammalian red blood cells (RBCs) relies upon the membrane skeleton, a viscoelastic network of short, membrane-associated actin filaments (F-actin) cross-linked by long, flexible spectrin tetramers. Nonmuscle myosin II (NMII) motors exert force on diverse F-actin networks to control cell shapes, but a function for NMII contractility in the 2D spectrin-F-actin network in RBCs has not been tested. Here, we show that RBCs contain membrane skeleton-associated NMIIA puncta, identified as bipolar filaments by super-resolution fluorescence microscopy. NMIIA association with the membrane skeleton is ATP-dependent, consistent with NMIIA motor domains binding to membrane skeleton F-actin and contributing to membrane mechanical stability. In addition, the NMIIA heavy and light chains are phosphorylated *in vivo* in RBCs, indicating active regulation of NMIIA motor activity and filament assembly, while reduced heavy chain phosphorylation of membrane skeleton-associated NMIIA indicates assembly of stable filaments at the membrane. Treatment of RBCs with blebbistatin, an inhibitor of NMII motor activity, decreases the number of NMIIA filaments associated with the membrane and enhances local, nanoscale membrane oscillations, suggesting decreased membrane tension. Blebbistatin-treated RBCs also exhibit elongated shapes, loss of membrane curvature, and enhanced deformability, indicating a role for NMIIA contractility in promoting membrane stiffness and maintaining RBC biconcave disc cell shape. As structures similar to the RBC membrane skeleton are conserved in many metazoan cell types, these data demonstrate a general function for NMII in controlling specialized membrane morphology and mechanical properties through contractile interactions with short F-actin in spectrin-F-actin networks.

**Significance statement:** The biconcave disc shape and deformability of the mammalian RBC is vital to its circulatory function, relying upon a 2D viscoelastic spectrin-F-actin network attached to the membrane. A role for myosin II (NMII) contractility in generating tension in this network and controlling RBC shape has never been tested. We show that NMIIA forms phosphorylated bipolar filaments in RBCs, which associate with F-actin at the membrane. NMIIA motor activity is required for interactions with the spectrin-F-actin network, and regulates RBC biconcave shape and deformability. These results provide a novel mechanism for actomyosin force generation at the plasma membrane, and may be applicable to other cell types such as neurons and polarized epithelial cells with a spectrin-F-actin-based membrane skeleton.

## Introduction

RBC biconcave disc shape and deformability are vital to circulatory function, providing a maximal surface-area-to-volume for optimal gas and ion exchange, and enabling repeated transit through narrow blood vessels less than half their diameter during the ~120 day RBC lifespan (1–3). Appropriate levels of RBC deformation also regulate blood flow and oxygen delivery via mechanotransductive pathways that release ATP to induce vasodilation and hyperemia (4). These properties all rely upon the membrane skeleton, a 2-dimensional (2D) quasi-hexagonal network of short (~37 nm) actin filament (F-actin) nodes interconnected by ~200 nm long, flexible (α_1_β_1_)_2_–spectrin tetramers (5, 6). Molecular genetics, biochemistry, biophysics and physiology of human and mouse congenital hemolytic anemias have shown that multiple proteins which mediate the horizontal connectivity of the micron-scale 2D network, along with the network’s multipoint vertical attachments to the membrane, are critical for RBC shape and deformability (1, 2, 5, 7). Similar membrane skeleton networks are a conserved feature of metazoan cells, where they create specialized membrane domains of ion channels, pumps, or cell adhesion molecules that confer complex signaling, cell interactions and mechanical resilience on the plasma membrane (8–10).

Since the first observation of the biconcave disc shape of RBCs nearly 200 years ago (11), a wealth of knowledge has accumulated regarding the molecular interactions of the membrane skeleton components and their contributions to RBC membrane properties and physiology. However, numerous unanswered questions remain regarding the forces behind the genesis and maintenance of RBC biconcave shape (5, 12, 13). Non-muscle myosin II, a force-generating motor protein, was identified and purified from RBCs >30 years ago and forms typical heterohexamers (termed NMII molecules) of two heavy chains (HCs), two regulatory light chains (RLCs), and two essential light chains (ELCs) (14, 15). RBC NMII also has a typical F-actin-activated Mg^++^ATPase activity that depends on RLC phosphorylation by myosin light chain kinase (MLCK) and assembles into bipolar filaments *in vitro*, similar to NMIIs from other cells (14–16). Mature human RBCs contain ~6,200 NMII molecules and ~500,000 actin molecules per RBC, resulting in about 80 actin molecules per NMII molecule, which is similar to that of other cells, such as platelets (15, 17). NMII in mature RBCs has been variously hypothesized to control RBC shapes (15, 18, 19), repair local disruptions in the spectrin-F-actin network (20), or to be retained as a non-functional vestige from an earlier, more motile stage of erythroblast terminal differentiation and maturation (17), but none of these hypotheses have been experimentally tested.

In most nucleated eukaryotic cells, NMII bipolar filaments generate tension to control membrane deformations and cell shape by pulling on F-actin in the cortex, a 3D network adjacent to the plasma membrane (21–24). However, cortical F-actin is longer (~100-600nm) than the filaments in the 2D spectrin-F-actin network (~37nm), and associates with a different complement of F-actin binding proteins (25). Thus, it is an open question whether NMII could pull on the short F-actins of the RBC membrane skeleton to generate tension and influence membrane properties such as shape, curvature, and mechanics. Because spectrin-F-actin networks are a conserved feature of metazoan cells, findings in RBCs could be broadly significant for elucidating fundamental principles of cytoskeletal organization and regulation of specialized plasma membrane structure and function. Since RBC F-actin is present exclusively in the spectrin-F-actin network of the membrane skeleton (5, 6), RBCs are an ideal model system to test this question, without confounding contributions from other cytoskeletal populations of F-actin.

In this study, we provide evidence that NMIIA is the predominant RBC NMII isoform, and that NMIIA is located in close proximity to the plasma membrane, where it is associated with F-actin in the membrane skeleton. Super-resolution and TIRF microscopy indicates that NMIIA can form bipolar filaments in intact RBCs, and that some of these filaments are associated with the membrane. Moreover, the RBC NMIIA HC and RLC are phosphorylated *in vivo*, indicating active regulation of NMIIA motor activity and filament assembly. Inhibition of NMIIA contractility with blebbistatin results in reduced NMIIA association with the membrane, RBC elongation and loss of biconcavity, with increased local and global membrane deformability in flickering and microfluidics assays. These results demonstrate a previously unrealized motor-dependent interaction between NMIIA and the spectrin-F-actin network that controls RBC membrane tension, biconcave disc shape and deformability.

## Results

### NMIIA, the predominant NMII isoform in human RBCs, forms bipolar filaments at the RBC membrane

The majority of cell types express more than one isoform of the NMII HC, typically the NMIIA HC encoded by the *MYH9* gene and the NMIIB HC encoded by the *MYH10* gene (22–24). A third NMIIC HC isoform, encoded by the *MYH14* gene, is also present in some cell types but is not as widely expressed as the *MYH9* and *MYH10* genes (26). To assess NMII isoform content during human erythroid differentiation, we measured mRNA and protein expression levels of NMIIA HC and NMIIB HC in human CD34^+^ cells induced to differentiate into erythroid cells *in vitro* (Fig. S1A). qRT-PCR shows that *MYH9* and *MYH10* transcripts are present during early stages in culture and decrease during the final stages of terminal erythroid maturation (Fig. S1B). Immunoblots of the same samples show that NMIIB HC protein decreases below detectable levels by day 10 of differentiation, while NMIIA HC protein persists through day 16 of culture (Fig. S1C), when most erythroblasts have extruded their nuclei and become reticulocytes (Fig. S1A). Moreover, immunoblotting of human RBC membranes (ghosts) isolated by hypotonic hemolysis and washing further indicates that NMIIA is the predominant NMII isoform in mature human RBCs (Fig. S1D). These data agree with previous 2D chymotryptic peptide maps of purified RBC NMII HC, which are nearly identical to those of platelet NMIIA HC (15). Taken together, the data suggest that NMIIA is the predominant NMII isoform present in human RBCs.

To characterize the distribution of NMIIA in RBCs, fixed biconcave RBCs were immunostained for the NMIIA motor domain and phalloidin-stained for F-actin. Due to the low abundance of NMIIA in RBCs (~6,200 molecules/cell; (15, 17)), and small sizes of RBCs (8μm diameter × 2μm thick; (27)), RBCs were imaged using sensitive super-resolution AiryScan confocal fluorescence microscopy, with a lateral resolution of ~140nm and an axial resolution of ~400nm (28). 3D reconstructions of Z-stacks revealed that NMIIA is present in puncta uniformly distributed throughout each RBC, distinct from the more continuous F-actin localization at the membrane (Fig. 1A). Quantification of the number of NMIIA puncta in each cell demonstrated ~150 puncta/RBC (Fig. 1D). To better evaluate the distribution of NMIIA puncta, we examined single optical sections of individual RBCs (Fig. 1B). NMIIA motor domain puncta are present throughout the RBC and at the edge of the RBC where they localize with F-actin at the membrane (Fig. 1B, arrowheads). Closely spaced doublets of NMIIA motor domain puncta were also identified spaced 200-400nm apart (Fig. 1B, arrows), equal to the lengths of NMIIA bipolar filaments observed *in vitro* and in cells using electron microscopy and super-resolution microscopy (29–32). To obtain unbiased data on the distances between all individual NMIIA motor domain puncta in RBCs, we performed 3D nearest-neighbor analysis (Fig. 1C), which showed that 49% of NMIIA motor domain puncta are within 200-400nm of one other puncta, forming doublet pairs that could represent NMIIA bipolar filaments (Fig. 1E). NMIIA puncta that are not within 200-400nm of one another may represent small filaments below the resolution of AiryScan, or other types of NMIIA assemblies.

**Figure 1.**
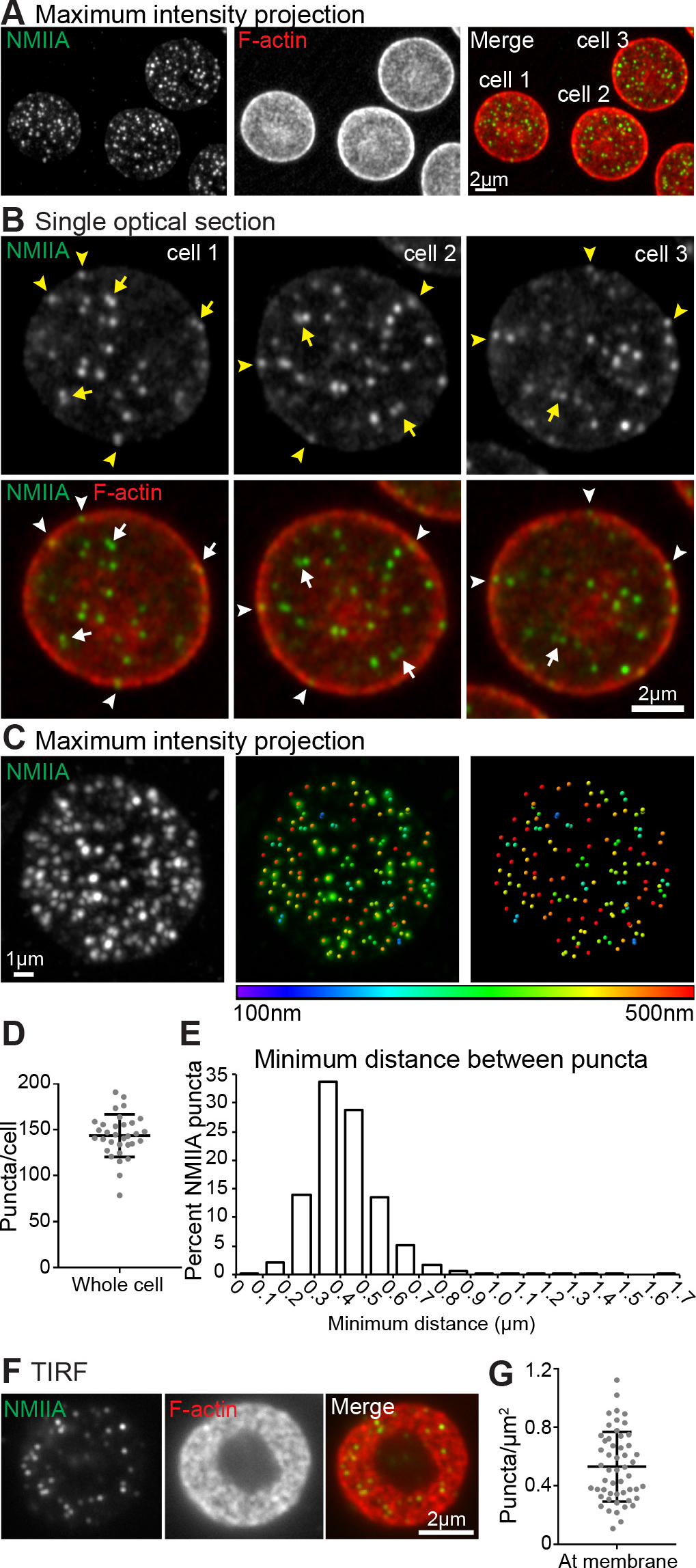
NMIIA localizes as uniformly distributed puncta in RBCs, likely representing bipolar filaments associated with the membrane. (A) Maximum-intensity projections of super-resolution AiryScan confocal Z-stacks of RBCs immunostained with an antibody to the motor domain of NMIIA (green) and rhodamine-phalloidin for F-actin (red). (B) Higher-magnification views of NMIIA motor domain puncta in single XY optical sections from the super-resolution AiryScan Z-stack shown in (A). (C) Maximum-intensity projection of a representative RBC stained for NMIIA (left). Center image shows NMIIA puncta computationally identified in Imaris (colored dots) superimposed on the NMIIA staining (green). Right image shows identified NMIIA puncta alone. Identified puncta are color-coded based on the minimum distance to the nearest puncta in 3D, with purple representing puncta ~100nm apart and red identifying puncta 500nm or more apart. (D) Quantification of the number of NMIIA puncta per cell from AiryScan images. Numbers were calculated by dividing the total number of puncta in each image by the number of cells in each image. n = 32 images (176 total cells from 2 individual donors). Mean ± S.D. = 143 ± 23 puncta/cell. (E) Histogram showing distribution of the minimum distances between NMIIA motor domain puncta measured from AiryScan images of whole RBCs similar to that in (A-C). n = 25067 puncta from 176 cells from 2 individual donors. Mean ± S.D. = 0.42 ± 0.13 μm. 49% of puncta are between 200-400 nm apart. (F) TIRF microscopy images of immunostaining for NMIIA motor domain (green) and rhodamine phalloidin for F-actin (red) in RBCs showing an *en face* view of the membrane surface within 100-200 nm of the coverslip. (G) Quantification of the density of NMIIA puncta/μm^2^ within 200 nm of the membrane from TIRF images. n = 52 RBCs from 2 individual donors. Lines represent mean ± S.D. (0.53 ± 0.24 puncta/μm^2^).

To better examine NMIIA motor domain association with F-actin at the membrane, we performed TIRF microscopy, which selectively illuminates fluorophores located within 100-200nm from the coverslip, at or near the plasma membrane (33). TIRF images of F-actin in biconcave RBCs reveal a donut-shaped appearance where the membrane of the rim is in close contact with the coverslip (Fig. 1F, Fig. S2A). Discrete NMIIA puncta appear scattered throughout this donut-shaped region, localizing with F-actin at the membrane (Fig. 1F, Fig. S2A), and are not observed in secondary antibody-alone controls (Fig. S2A). Quantification indicates an average of ~0.5 puncta/μm^2^ in TIRF images of biconcave RBCs (Fig. 1G). Similar patterns and numbers of NMIIA puncta at the membrane are also observed by epifluorescence and TIRF microscopy of RBCs stained with antibodies to the NMIIA tail domain (Fig. S2B-D), similar to previous epifluorescence microsocopy using polyclonal antibodies to NMIIA (15). Based on a total RBC surface area of 140μm^2^ (34), these data show that ~70 NMIIA puncta are associated with the membrane of each RBC.

### NMIIA association with the Triton X-100-insoluble membrane skeleton is ATP-dependent

A contractile function for NMIIA bipolar filaments in RBCs would be expected to involve MgATP-dependent interactions of NMIIA motor domains with F-actin in the membrane skeleton (22–24). The RBC membrane skeleton is defined operationally as the insoluble network of spectrin, actin and associated proteins remaining after non-ionic detergent extraction of membrane lipids and integral membrane proteins (35). Therefore, purified RBC membranes (Mg^++^ ghosts) containing NMIIA were prepared by hypotonic hemolysis (15), extracted with Triton X-100 (TX-100) to solubilize lipids and integral membrane proteins, sedimented, and subjected to immunoblot analysis for NMIIA HC (Fig. 2A, B). Note that the decreased mobility for the NMII HC in the ghosts and the TX-100-extracted pellet samples on these acrylamide gradient mini-gels is due to the large excess of spectrin over NMIIA HC which forces the HC to migrate faster, and is not observed on large format, low percentage gels (Figure S3). Approximately 40% of the NMIIA HC in Mg^++^ ghosts is associated with the TX-100-insoluble membrane skeleton in the absence of MgATP (Fig. 2A, C), decreasing to 10% in the presence of MgATP (Fig. 2B, C). In contrast, nearly all the actin is associated with the membrane skeleton in both the absence and presence of ATP (Fig. 2A-C). All of the spectrin, actin, and other membrane skeleton proteins are present in the TX-100-insoluble pellet in the absence or presence of MgATP, while the majority of the major integral membrane protein, Band 3, is in the supernatant, as expected (Fig. S3A). Thus, the MgATP-dependent decrease in NMIIA associated with the membrane skeleton is not due to selective extraction of actin or other membrane skeleton proteins. In agreement with these results, when intact RBCs are extracted with TX-100 in the presence of endogenous ATP, little NMIIA remained associated with the TX-100-insoluble pellet (Fig. S3B, C). By contrast, ~55% of NMIIA is associated with the TX-100 insoluble pellet when cellular ATP levels are depleted by incubation of the RBC lysate with hexokinase and glucose prior to sedimentation (Fig. S3B-C). These data show that NMIIA associates with the membrane skeleton via MgATP-dependent binding to F-actin, consistent with *in vitro* biochemical and biophysical studies showing that MgATP weakens purified NMII motor domain binding to F-actin (36).

**Figure 2.**
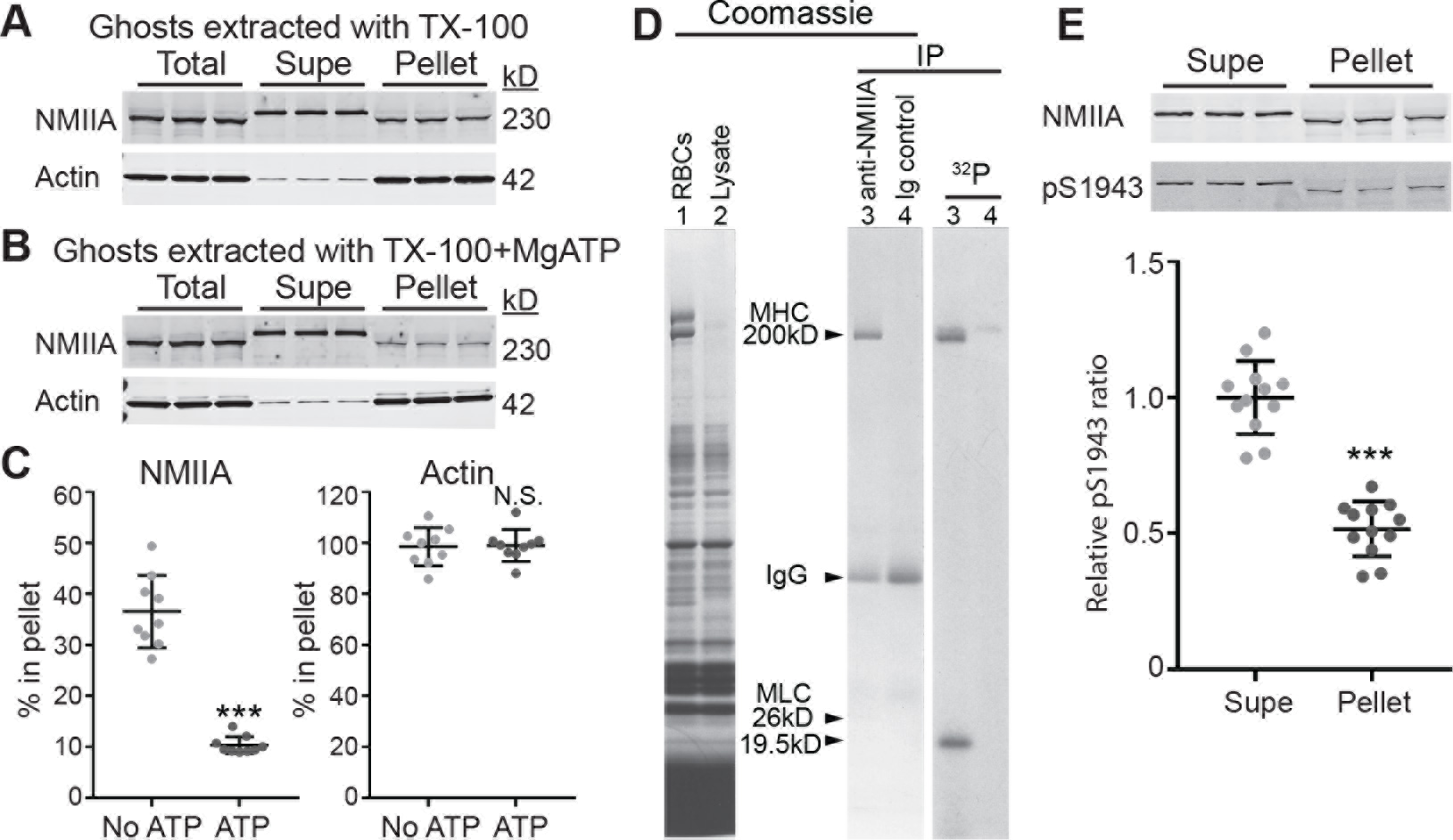
NMIIA association with the RBC membrane skeleton is MgATP dependent, and NMIIA HC and RLC phosphorylation indicates active regulation of NMIIA motor activity and filament assembly at the membrane skeleton. (A-B) RBC Mg^++^ ghosts were extracted in TX-100 buffer without (A) or with (B) addition of 5mM MgATP followed by SDS-PAGE and immunoblotting for NMIIA HC or actin in the soluble (Supe) and insoluble membrane skeleton (Pellet) fractions. (C) Quantification of % NMIIA or actin in the membrane skeleton fraction. 36.5 ± 7.1% NMIIA is in the TX-100-insoluble pellet in the absence of MgATP, and 10.4 ± 1.7% is in the pellet in the presence of MgATP, p < 0.0001. 98.5 ± 7.5% actin is in the TX-100-insolubule pellet with no ATP, and 98.9 ± 6.25% is in the pellet with ATP. Each point in C represents one of three technical replicates for each of three biological replicates (ghosts from 3 donors), for a total n = 9. Lines represent mean ± S.D. (D) Live RBCs were metabolically labeled with ^32^P-orthophosphate to detect phosphorylated proteins, lysed, and native NMIIA was immunoprecipitated (IP) under non-denaturing conditions. Left panel, Coomassie blue gel of total RBCs (lane 1) and Lysate before IP (lane 2). Right panel, Coomassie blue gel of anti-NMIIA IP (lane 3) and preimmune IgG IP (lane 4), and autoradiogram of this gel. The RBC NMIIA 26kD and 19.5kD MLC were visible in the original Coomassie-stained gel, but faded upon destaining (15). (E) RBC Mg^++^ ghosts were extracted in TX-100 buffer followed by SDS-PAGE and immunoblotting for NMIIA HC (top) or NMIIA HC phosphorylated on serine 1943 (pS1943) (bottom). Quantification shows that the ratio of pS1943 HC/total HC is about two times higher in the TX-100-soluble supernatant (mean ± S.D. = 1.0 ± 0.13) compared to the TX-100-insoluble pellet (mean ± S.D. = 0.52 ± 0.10). Each point represents one of three technical replicates for each of four biological replicates (4 individual donors), for a total n = 12. Lines represent mean ± S.D. p < 0.0001.

### Phosphorylation of NMIIA HC and RLC is consistent with contractility and regulated filament assembly in the membrane skeleton

NMII motor activity and filament assembly are regulated by signaling pathways that result in the phosphorylation of multiple sites near the N-terminus of the RLC and near the C-terminus of the HC (22–24, 37). To determine if RBC NMIIA HC or RLC is phosphorylated *in vivo*, RBCs were metabolically labeled with ^32^P orthophosphate. Immunoprecipitation of native NMIIA under conditions that preserve HC and LC associations followed by SDS-PAGE and autoradiography reveals that both the 19.5-kDa RLC and the HC are phosphorylated, suggesting active regulation of NMIIA motor activity and filament assembly in live RBCs (Fig. 2D). To test whether NMIIA HC phosphorylation status affects association with the membrane skeleton, we extracted Mg^++^ ghosts in TX-100 as described above and immunoblotted for total NMIIA HC and NMIIA HC phosphorylated at serine 1943. The ratio of phosphorylated HC to total HC was about two-fold greater in the TX-100-soluble supernatant as compared to the TX-100 insoluble pellet containing the membrane skeleton (Fig. 2E). As HC phosphorylation at serine 1943 inhibits NMIIA filament assembly and promotes filament turnover (37), these data suggest that the NMIIA associated with F-actin in the membrane skeleton consists of NMIIA filaments with reduced HC phosphorylation and increased stability.

### NMIIA motor activity affects NMIIA association with the RBC membrane

Functional interactions of NMIIA filaments with F-actin in the RBC membrane skeleton would be expected to depend on motor domain activity. To evaluate whether this influences distribution of NMIIA and F-actin at the RBC membrane, we treated live, intact RBCs with blebbistatin, an NMII-specific motor inhibitor that prevents phosphate release and stalls NMII in a weak F-actin-binding state, blocking contractility (36). As negative controls for the effects of blebbistatin, we also treated RBCs with DMSO or the inactive enantiomer of blebbistatin. NMIIA association with the membrane was evaluated by TIRF microscopy of fixed RBCs immunostained for the NMIIA motor domain and flattened by cytospinning, providing a greater membrane surface area to examine (Fig. 3A). Blebbistatin treatment of RBCs resulted in a ~30% reduction in NMIIA puncta/μm^2^ at the membrane relative to DMSO treatment, while the inactive enantiomer of blebbistatin had no significant effect (Fig. 3B). The decrease in NMIIA puncta density at the membrane was not due to changes in the extent of RBC flattening prior to imaging, since blebbistatin had no effect on the area of RBC F-actin visualized by TIRF microscopy (Fig. 3C). Thus, the localization and distribution of NMIIA at the RBC membrane partially depend on NMIIA motor activity, supporting a contractile interaction of NMIIA motor domains with F-actin in the membrane skeleton.

**Figure 3.**
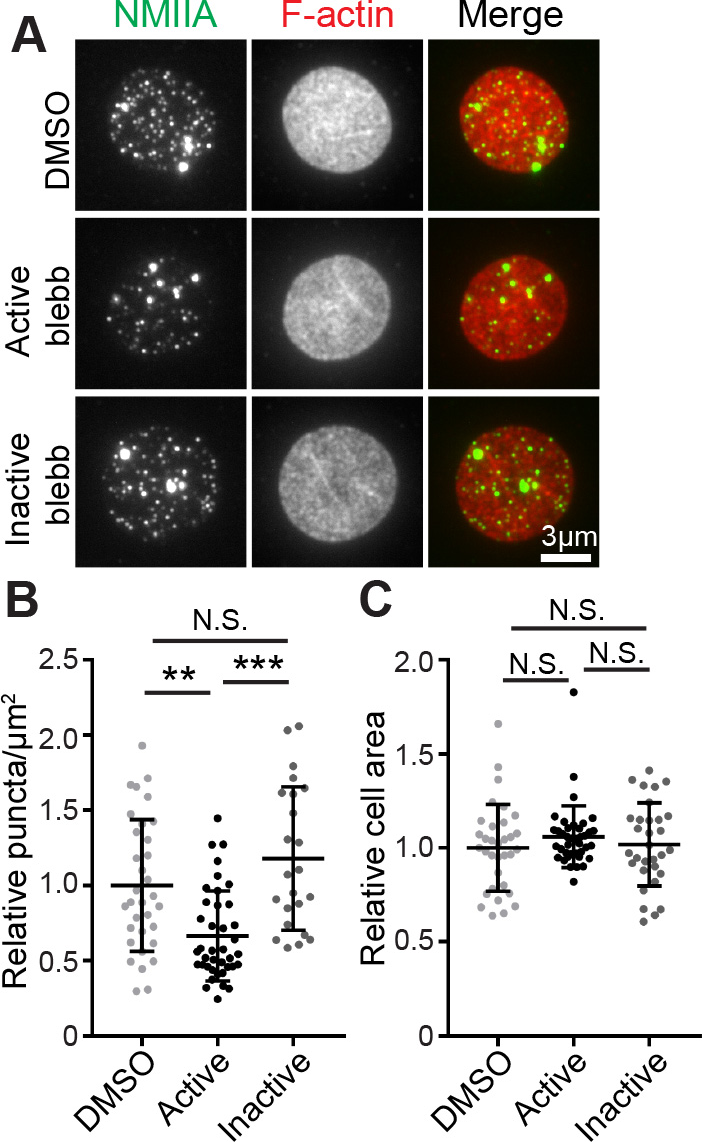
NMIIA motor activity controls the association of NMIIA puncta with the RBC membrane. (A) TIRF microscopy of NMIIA motor domain (green) and rhodamine-phalloidin for F-actin (red) in RBCs pre-treated with DMSO alone, 20μM active blebbistatin, or 20μM inactive blebbistatin before fixation and immunostaining. RBCs were flattened prior to imaging to visualize a larger area of the membrane. (B) NMIIA puncta density at the RBC membrane measured as the number of NMIIA puncta per μm^2^ in TIRF images. DMSO versus active blebbistatin p = 0.0016, active blebbistatin versus inactive blebbistatin p < 0.0001. (C) Cell surface areas for RBCs in TIRF images in each treatment group. There is no significant difference in cell areas between treatment groups by one-way ANOVA. DMSO, n = 33; active blebbistatin, n = 40; inactive blebbistatin, n = 31 cells from 2 individual donors.

To determine whether inhibition of NMIIA motor activity affected F-actin organization in the membrane skeleton, we examined the TIRF images of phalloidin-stained RBCs treated with DMSO, active blebbistatin, or inactive blebbistatin. In all three treatment groups, the F-actin appeared as a dense reticular network, and no detectable differences in F-actin staining were apparent in active blebbistatin-treated RBCs as compared to controls (Fig. S4A). In addition, immunoblotting of RBC cytosol for actin showed that blebbistatin treatment did not lead to increased G-actin in the cytosol due to F-actin depolymerization from the membrane skeleton (Fig. S4B, C). Together, these data indicate inhibition of NMIIA motor activity in RBCs does not lead to global rearrangements or disassembly of the spectrin-F-actin network.

### NMIIA motor activity controls RBC morphology and membrane deformability

Appropriate levels of membrane tension, provided by the spectrin-F-actin network attached to transmembrane proteins, are required for the maintenance of RBC biconcave shape (1, 2, 7). Because treatment with blebbistatin altered NMIIA association with the membrane, we predicted that blebbistatin inhibition of NMIIA motor activity would lead to a loss in membrane tension, leading to changes in RBC membrane curvature and shape. To test this hypothesis, we treated RBCs with DMSO, active blebbistatin, and inactive blebbistatin as above. Confocal fluorescence microscopy of RBCs immunostained for glycophorin A (GPA), a membrane marker (Fig. 4A), revealed that treatment with 20μM active blebbistatin caused RBCs to elongate, measured as an increased aspect ratio (major/minor axis) in XY maximum intensity projections (Fig. 4B). In addition, blebbistatin treatment led to a loss of biconcavity as measured by an increased height ratio (minimum/maximum height) of XZ slices from the center of each RBC (Fig. 4C). Neither DMSO nor inactive blebbistatin induced significant RBC elongation or loss of biconcavity (Fig. 4A-C), and increasing the concentration of active blebbistatin to 50 μM did not lead to further increases in RBC elongation or loss of biconcavity (Fig. S5). Interestingly, there was a positive relationship between RBC elongation and loss of biconcavity in RBCs from all treatment groups, although the high variability of the data reduced the extent of the correlation (Fig. S6A). In addition, we found that changes in biconcavity arose within 30 minutes of treatment with active blebbistatin, while changes in the aspect ratio did not arise until two hours of treatment (Fig. S6B-C). Thus, while both RBC elongation and biconcavity depend on NMIIA motor activity, these two parameters vary independently in many cells. These results indicate that NMIIA motor activity is necessary for the maintenance of RBC membrane curvature characteristic of the biconcave disc shape.

**Figure 4.**
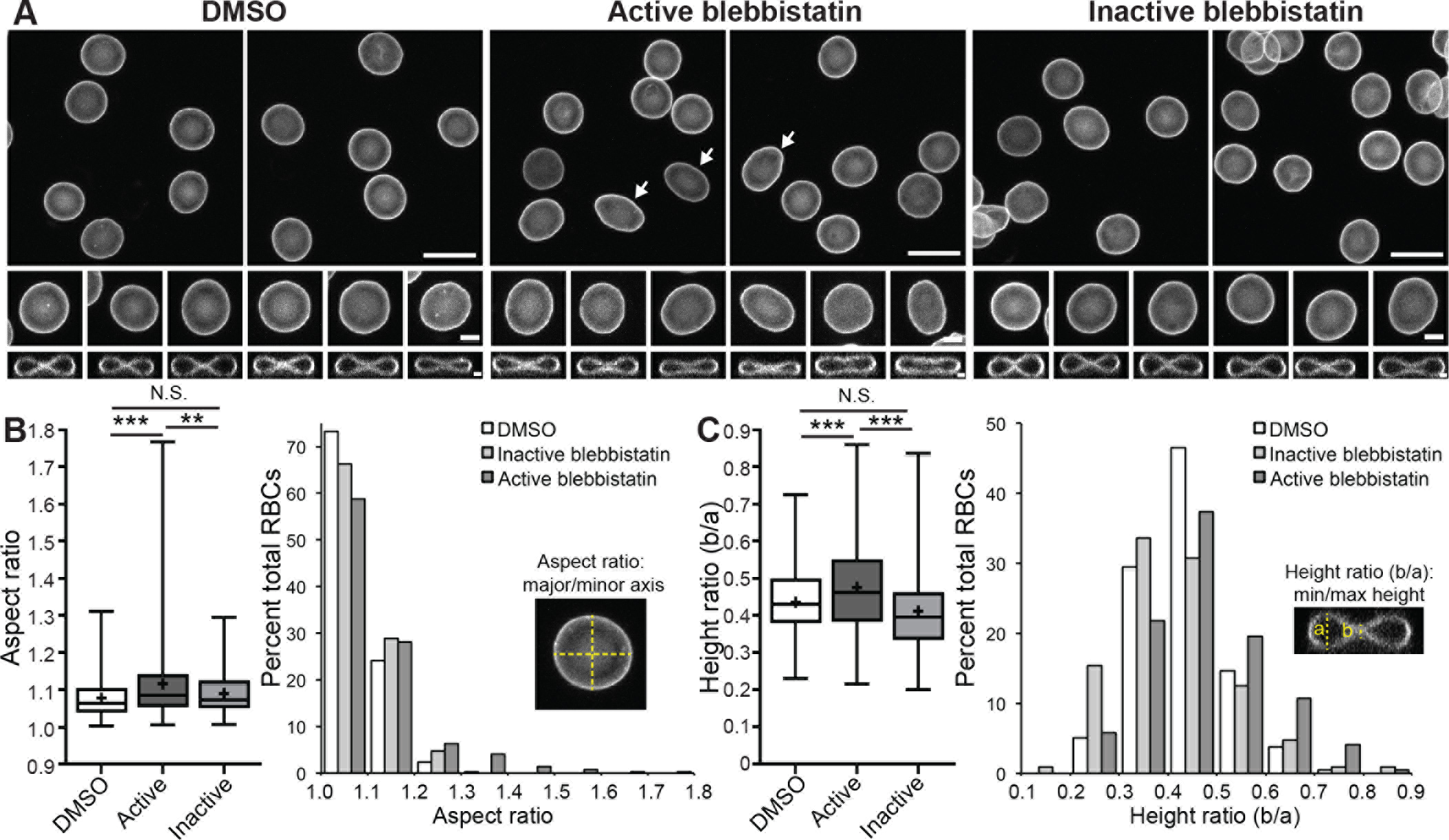
NMIIA motor activity controls RBC morphology. (A) Confocal fluorescence microscopy images of GPA-stained RBCs after treatment with DMSO, 20μM active blebbistatin, or 20μM inactive blebbistatin. Top row (scale = 10μm) and middle rows (scale = 2μm) show XY maximum-intensity projections, while the bottom row (scale = 1μm) shows XZ slices from the middle of the cell. DMSO- and inactive blebbistatin-treated control RBCs are round in XY and biconcave in XZ views, as expected. Note elongated cells and cells with relaxed dimples after treatment with active blebbistatin (arrows). (B) RBC elongation measured from aspect ratios (major/minor axis) in XY maximum-intensity projections as in A. Left, box-and-whisker plot; right, histogram. DMSO versus active blebbistatin, p < 0.0001. Active versus inactive blebbistatin, p = 0.0071. (C) RBC biconcavity measured from the ratio of cell height at the dimple to height at the rim from XZ slices as in A. Left, box-and-whisker plot; right, histogram. DMSO versus active blebbistatin, p < 0.0001. Active versus inactive blebbistatin, p < 0.0001. For box-and-whisker plots, middle line represents median, upper and lower lines represent third and first quartiles respectively, and whiskers represent minimum and maximum values. + sign denotes means. DMSO, n = 423 RBCs (4 donors); Active blebbistatin, n = 410 RBCs (4 donors); inactive blebbistatin, n = 104 RBCs (1 donor).

Maintenance of RBC membrane curvature could be due to NMIIA contractility promoting membrane tension and reducing local membrane deformability. A sensitive measure of local membrane deformability is the spontaneous, nanoscale membrane oscillation observed by changes in light scattering at the RBC surface using positive low phase-contrast video microscopy (38–40). To examine the role of NMIIA activity in these oscillations, we filmed live RBCs treated with DMSO or active blebbistatin. Computational analysis of the amplitudes of RBC membrane oscillations showed that blebbistatin disruption of NMIIA activity increases flickering amplitudes (Fig. 5A, B), as well as their coefficient of variation (Fig. 5C, D). Thus, NMIIA motor activity appears to dampen spontaneous membrane oscillations in untreated RBCs, suggesting NMIIA contractility maintains membrane tension and reduces local deformability of the RBC membrane.

**Figure 5.**
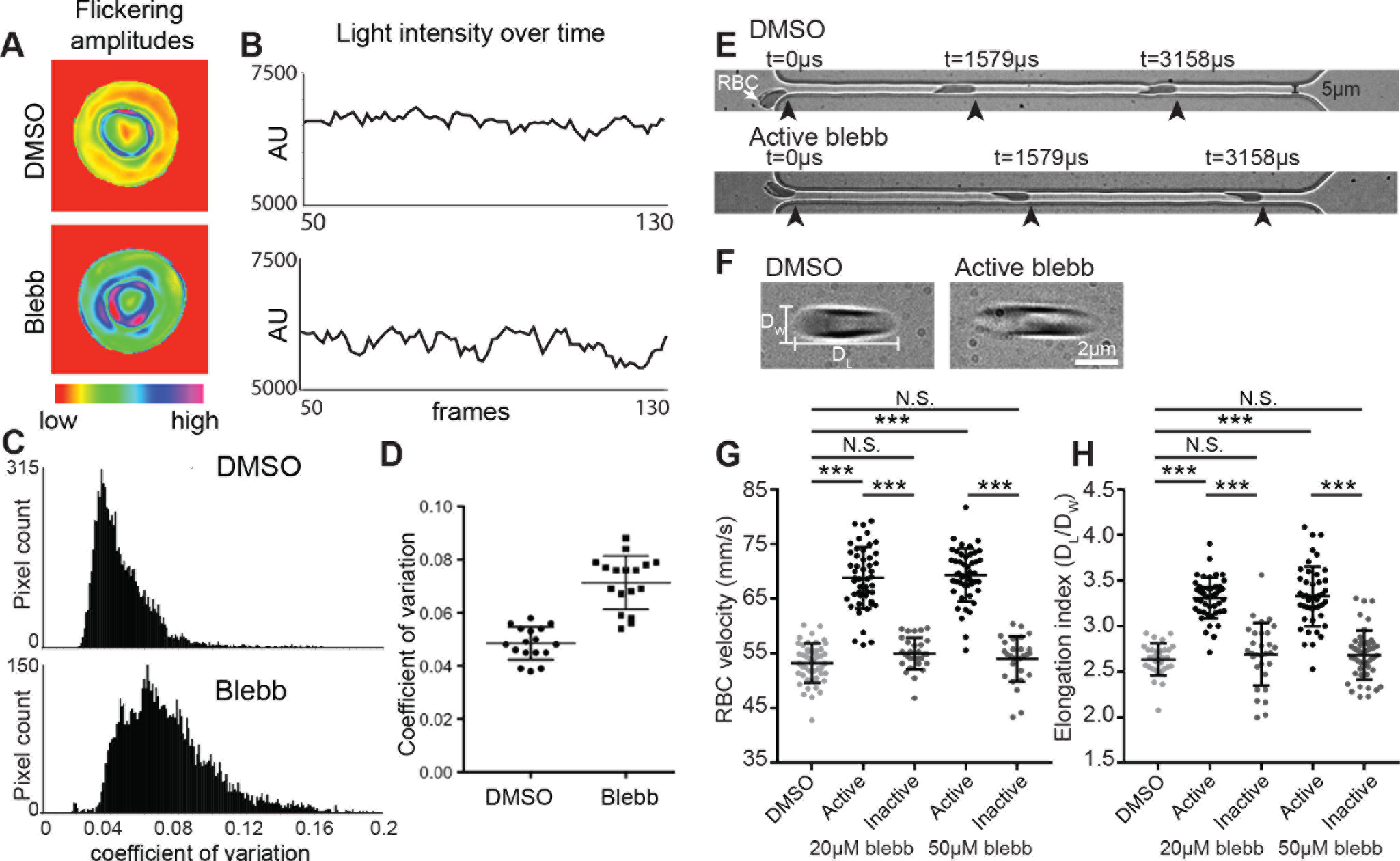
NMIIA motor activity controls the amplitude and variability of RBC membrane flickering and RBC deformability during flow in microchannels. (A-D) RBCs were treated with 5μM active blebbistatin or DMSO. (A) Color-coded representations of whole-RBC flickering amplitudes in representative cells, where red represents amplitude changes near zero and purple represents maximum changes. The blebbistatin-treated cell has higher flickering amplitudes throughout the cell. (B) Changes in light intensity over time in an area of 3×3 pixels in representative cells, as recorded by phase-contrast microscopy. The area from the blebbistatin-treated cell has larger changes in light intensity. (C) Frequency distributions of the coefficients of variation in all pixel amplitudes from representative cells. The blebbistatin-treated cell has more pixels with high coefficients of variation. (D) Coefficients of variation of membrane flickering amplitudes. n = 15 RBCs per treatment condition. (E, G) Treatment with active blebbistatin, but not inactive blebbistatin or DMSO, increases the velocity of RBCs flowing through a 5μm-wide channel, corresponding to an increase in deformability. Representative cells are shown in (E). In (G), DMSO n = 48; 20 μM active blebbistatin n = 48; 20 μM inactive blebbistatin n = 30; 50 μM active blebbistatin n = 47; 50μM inactive blebbistatin n = 28. (***, p < 0.0001). Data are presented as mean ± S.D. for each condition. (F, H) Treatment with active, but not inactive blebbistatin or DMSO increases the shear-induced elongation index as RBCs flow through a 20μm-wide constriction in a 30μm-wide channel, corresponding to an increase in deformability. Representative cells are shown in (F). In (H), DMSO n = 30; 20 μM active blebbistatin n = 45; 20 μM inactive blebbistatin n = 30; 50 μM active blebbistatin n = 45; 50 μM inactive blebbistatin n = 45. (***, p < 0.0001). Data are presented as mean ± S.D. for each condition.

RBC biconcave shape and local membrane deformability are important for whole-RBC deformability during RBC transit through small capillaries *in vivo* (2, 3). We evaluated RBC deformability *ex vivo* by measuring rates of RBC flow through a microfluidic device with channel dimensions similar to small capillaries, narrower than the 8μm diameter of human RBCs (Fig. 5E) (41). Using high-speed video microscopy we measured individual RBC velocities traveling through a narrow (5μm) channel and found that RBCs treated with 20μM active blebbistatin traveled through the channels more rapidly than DMSO- and inactive blebbistatin-treated control RBCs (Fig. 5G). An effect of a similar magnitude occurred in RBCs treated with 50μM blebbistatin (Fig. 5G). We also assessed the effect of blebbistatin on shear-induced deformation by measuring the elongation index of RBCs as they passed through a wide (20μm) constriction in a 30μm channel (Fig. 5F). This experiment showed that blebbistatin-treated RBCs became more elongated than control RBCs during flow through these wider channels (Fig. 5H). Again, an effect of a similar magnitude occurred in RBCs treated with 50μM blebbistatin (Fig. 5H). Blebbistatin treatment did not affect the responses of RBC flow through microchannels or RBC elongation in shear flow to changes in the oxygen pressure (PO_2_) (Fig. S7). As inhibition of NMIIA motor activity with blebbistatin increases whole-RBC deformability and ability to traverse microchannels, we conclude that NMIIA motor activity negatively regulates these properties in untreated RBCs. This is consistent with NMIIA promotion of membrane tension and maintenance of membrane curvature and biconcave shape.

## Discussion

The membrane skeleton, a long-range 2D periodic network of short F-actins cross-linked by flexible spectrin molecules, underlies the plasma membrane of metazoan cells, including RBCs. Here, we identified NMIIA contractility with the spectrin-F-actin network as a mechanism controlling the shape and membrane deformability of human RBCs. Fluorescence microscopy, as well as biochemical and biophysical approaches show that NMIIA bipolar filaments interact with the F-actin of the network via their motor domains to increase local and global membrane stiffness and maintain RBC biconcave disc shape. This may represent a previously unrecognized – and potentially conserved – paradigm for actomyosin regulation of membrane tension, curvature, cell shape, and biomechanics.

The NMIIA and NMIIB HC isoforms, encoded by the *MYH9* and *MYH10* genes, respectively, are both expressed in immature human erythroid progenitors, and NMIIB has been shown to play a role during terminal differentiation and enucleation (42). However, NMIIA persists into late stages of terminal erythroid maturation and in mature human RBCs, while NMIIB decreases below detectable levels. This data agrees with recent proteomics studies indicating that NMIIA and NMIIB levels decrease during terminal erythroid maturation, and that NMIIA is up to 10-fold more abundant than NMIIB in mature RBCs (43–45). While some proteomics studies identify NMIIB in mature RBCs, this apparent disagreement with our western blot data could be due to the increased sensitivity of mass spectrometry and/or contamination from leukocytes.

Super-resolution fluorescence microscopy and 3D image analysis reveal ~150 NMIIA puncta/RBC, likely representing NMIIA filaments or other assemblies, since individual NMIIA molecules would not be detected using this approach. This is in good agreement with a previous biochemical analysis that estimated NMIIA could form ~210 filaments/RBC, based on ~6,200 NMII monomers/RBC and assuming ~30 monomers/filament (15). Only ~50% of puncta are within 200-400 nm of their neighbors, the expected size of NMIIA bipolar filament lengths *in vitro* and in cells (29–32). The other, more distantly-spaced NMIIA puncta may represent immature NMIIA filaments shorter than ~140nm (the AiryScan XY resolution), or filaments oriented in the axial direction where the Z resolution (~300nm) is unable to resolve the two ends of shorter bipolar filaments. Alternatively, some of the brighter NMIIA puncta could represent higher-order associations between multiple filaments that cannot be resolved. NMIIA assembly into variably sized filaments, which then assemble into higher-order structures, is characteristic of cells that rely on NMIIA for contractility (31, 32, 46).

TIRF microscopy reveals that ~70 NMIIA puncta/RBC localize within 100-200nm of the membrane, along with the F-actin of the membrane skeleton. This is about half of the NMIIA puncta detected in 3D reconstructions of AiryScan images of whole RBCs, though the puncta may not represent identical structures due to differences in resolution between TIRF and AiryScan microscopy. Immunoblotting assays also show that ~40-50% of the NMIIA HC is associated with the Triton-insoluble membrane skeleton. This is in contrast to spectrin, actin, and other previously characterized components of the membrane skeleton, which are nearly completely associated with the Triton-insoluble membrane skeleton (35, 47). Notably, inclusion of MgATP reduces NMIIA co-sedimentation with the Triton-insoluble membrane skeleton several-fold, as originally observed for association of myosin II with F-actin at the membrane in *Dictyostelium discoideum* (48). This suggests that NMIIA interacts with the RBC membrane skeleton via motor domains binding to F-actin. Moreover, decreased numbers of NMIIA puncta are present at the membrane in RBCs treated with blebbistatin, which maintains motor domains in a weak F-actin-binding state. Together, these observations suggest that NMIIA filaments may associate with F-actin in the membrane skeleton via dynamic and transient interactions of their motor domains. NMIIA may be particularly suited for such hypothesized transient interactions, based on a greater F-actin-activated ATPase activity and a lower duty ratio compared to NMIIB (22–24, 29). Direct evaluation of this hypothesis would require live imaging of fluorescently tagged NMIIA and F-actin in RBCs, and remains technically challenging.

NMII contractility is regulated by RLC phosphorylation, which activates NMII motor activity and competence for filament assembly, and by HC phosphorylation, which enables dynamic turnover of filaments (22–24, 37). RBC NMIIA is phosphorylated on both the RLC and HC, suggesting at least some of the NMIIA in RBCs is constitutively active. The reduced HC phosphorylation at serine 1943 in the NMIIA associated with the Triton-insoluble membrane skeleton suggests that this population of NMIIA forms stable filaments, consistent with the NMIIA puncta visualized associated with the membrane in fixed RBCs. Interestingly, RLC phosphorylation inhibits binding of NMIIA HCs to anionic phospholipids *in vitro* (49), raising the possibility that inactive NMIIA (i.e., without a phosphorylated RLC) may bind to RBC membrane lipids independent of motor domain interactions with F-actin. RBCs contain Rho/Rho-kinase (ROCK) pathway components (50), and proteomic studies have also suggested that they contain myosin light chain kinase, casein kinase, and protein kinase C, which can phosphorylate either the RLC or the C-terminus of the HC (22–24, 37). However, the upstream signaling pathways that may activate or inhibit these kinases and regulate NMIIA contractility in RBCs are unknown.

In addition, other F-actin-binding proteins could modulate RBC NMIIA activity. Tropomyosin isoforms regulate the F-actin-activated MgATPase and F-actin sliding velocity of NMIIA and other myosin isoforms. RBCs contain two tropomyosins, Tpm3.1 and Tpm1.9, present in equimolar amounts and extending along the length of each short F-actin in the membrane skeleton (5, 51–53). RBCs also contain caldesmon (2 caldesmons/short F-actin), which, together with tropomyosin, regulates F-actin-activated myosin ATPase activity *in vitro* (5, 19). A Tpm3.1-deficient mouse in which RBCs have abnormal shapes and deformability (47, 54) will provide an opportunity to test whether Tpm3.1 regulates NMIIA in RBCs *in vivo*.

Blebbistatin inhibition of NMIIA motor activity leads to altered RBC shapes, including elongation and loss of biconcavity, indicating that constitutive NMIIA contractility maintains RBC biconcave disc shapes. The morphological changes in RBCs due to inhibition of NMIIA contractility are similar to, but not as pronounced as those seen in human and mouse RBCs with mutations in integral membrane skeleton proteins. In these cases, mutant RBCs with disruptions in the spectrin-F-actin network linkages can be highly elongated (elliptocytic), while mutant RBCs with loss of network attachments to the membrane are typically round with variable sizes due to loss of membrane surface area (spherocytic) (2, 55). This difference in phenotype is not necessarily unexpected, as our data indicate that NMIIA is not an integral membrane skeleton protein and may associate with the spectrin-F-actin network in a dynamic and transient manner.

In endothelial cells migrating in 3D collagen gels, NMII contractility is associated with regions of negative plasma membrane curvature (56), which is analogous to the central region (dimple) of the biconcave RBC. Although we do not detect selective enrichment of NMIIA in the dimple region (or at the rim), it is possible that activated NMIIA with phosphorylated RLC could be differentially localized to the dimple or rim. Alternatively, NMIIA contractility distributed across the entire membrane may maintain tension in the spectrin-F-actin network coupled to the membrane bilayer, promoting membrane flattening and dimple formation, according to theoretical models for RBC biconcave shape (2, 57). To date, these models have not considered an active role for myosin-based contractility in generating tension and controlling RBC shape.

In addition to NMIIA maintenance of RBC biconcave shape, NMIIA contractility reduces RBC membrane deformability. Blebbistatin inhibition of NMIIA motor activity increases the variation of the amplitude of RBC membrane flickering, indicating that NMIIA contractility normally dampens these local membrane deformations. Cycling of NMIIA motor activity on F-actin may also partly contribute to the MgATP-dependence of membrane flickering in untreated RBCs (38, 40). Moreover, NMIIA contractility controls global RBC deformability under physiological flow conditions. Blebbistatin treatment increased the velocity of RBCs traveling through a narrow (5μm) channel, indicative of increased RBC membrane deformability. In a related assay, blebbistatin also increased the elongation of RBCs traveling through a 20μm constriction in a 30μm-wide channel, consistent with increased deformability upon inhibition of NMIIA contractility. RBC membrane flickering and transit through narrow channels also depend on F-actin assembly (38, 58), suggesting NMIIA contractility and F-actin polymerization may synergistically modulate membrane mechanical properties in RBCs.

In conclusion, we propose that NMIIA forms bipolar filaments associated via their motor domains with the short F-actins in the spectrin-F-actin network underlying the RBC membrane (Fig. 6A). Contractile forces exerted by NMIIA bipolar filaments on the spectrin-F-actin network could promote membrane tension to maintain RBC biconcave disc shape and restrict membrane deformation under shear flow. In support of this proposal, experimental inhibition of NMIIA contractile activity with blebbistatin decreases the association of NMIIA with F-actin at the membrane, as indicated schematically in Figure 6B, disrupting the RBC biconcave disc shape and increasing RBC deformability. NMIIA bipolar filaments likely exert force on only a small subset of RBC F-actins, as NMIIA puncta have a low density at the membrane (0.5 filaments/μm^2^) as compared to ~250 F-actins/μm^2^ (5, 6). Also, blebbistatin-induced NMIIA dissociation from the membrane does not result in detectable rearrangements of F-actin by TIRF imaging. Therefore, we hypothesize that NMIIA exerts tension on the RBC membrane by acting as a dynamic cross-linker between short F-actins, which are constrained by their linkages in the network (Fig. 6A), rather than by inducing large-scale F-actin translocation and/or rearrangements of the spectrin-F-actin network. NMIIA/F-actin cross-linking activity may be important for the ability of the spectrin-F-actin network to enhance RBC resilience and preserve biconcave disc shapes to optimize circulatory function *in vivo*. Future studies will determine the importance of NMIIA and other NMII isoforms for the functions of spectrin-F-actin networks in membrane morphology and resilience in other cell types.

**Figure 6.**
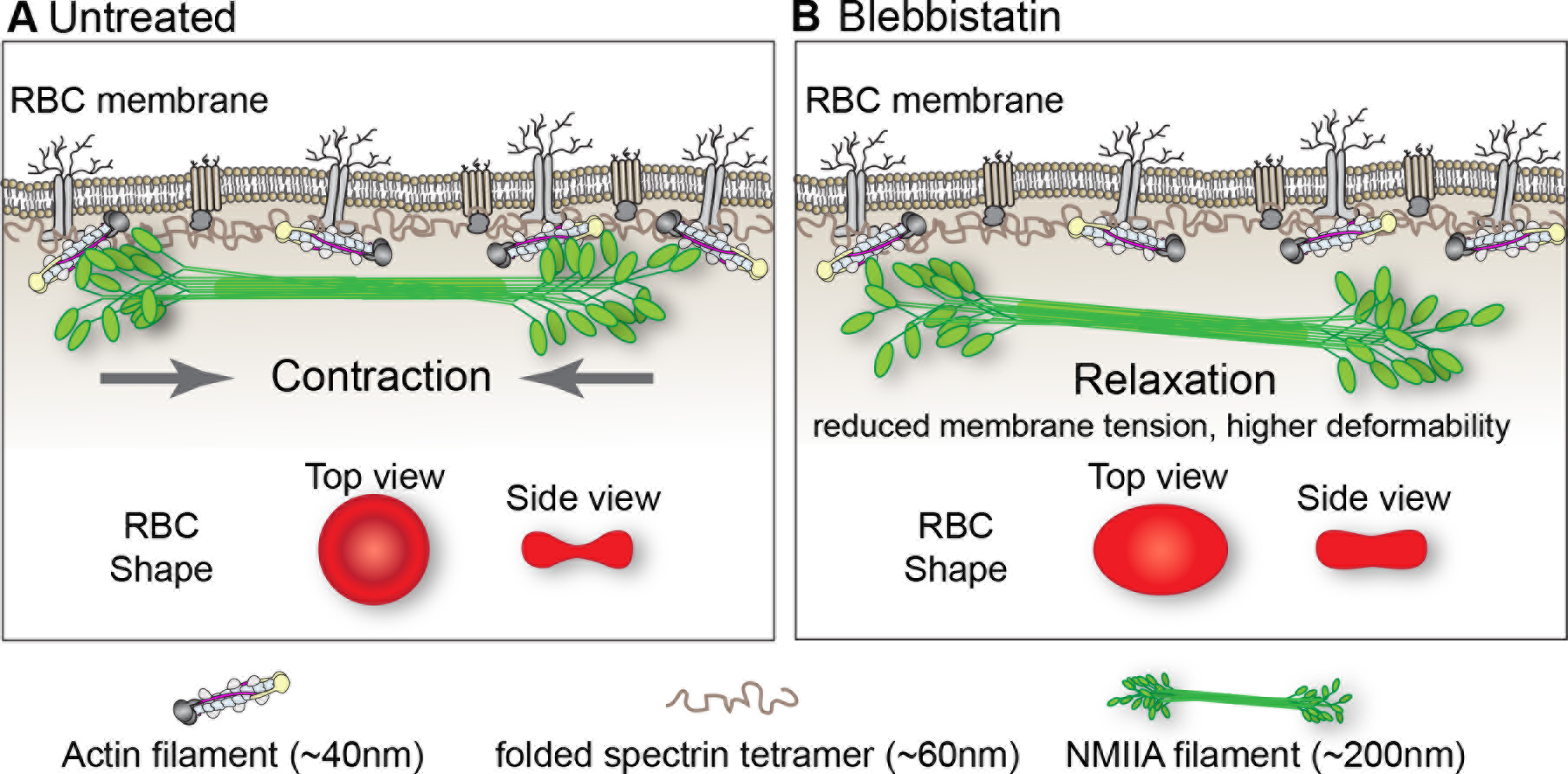
Model for NMIIA contractility in the spectrin-F-actin network and its effect on RBC membrane morphology and mechanical properties. (A) A 2D network of long, flexible spectrin tetramers that cross-link short actin filaments (the spectrin-F-actin network) is attached to the plasma membrane (not drawn to scale) via associations of network components with transmembrane proteins. NMIIA bipolar filaments can associate with the short F-actins of the spectrin-F-actin network via their motor domains. Through these interactions, NMIIA contractile forces (arrows) could promote membrane tension in the spectrin-F-actin network to maintain normal biconcave RBC shapes, shown in top and side views (red RBCs), and control RBC deformability. The NMIIA filament depicted in this diagram is ~200nm long, but our data and other studies show that myosin filaments can be up to 400nm long in cells. This would double the length scale over which one NMIIA filament could generate contractile forces on the membrane. (B) Blebbistatin treatment weakens the association between NMIIA motor domains and F-actin, causing NMIIA filaments to partially or completely dissociate from the spectrin-F-actin network. This dissociation would lead to a relaxation of the contractile forces on the network, reducing membrane tension and increasing deformability. This relaxation leads to an elongation of RBC shape, shown in the top view, and a decrease of biconcavity, shown in the side view (red RBCs). As indicated in the legend at the bottom of the figure, the actin filament barbed ends are capped by adducin (yellow) and their pointed ends by tropomodulin (dark gray), with tropomyosin rods (purple) spanning their length, and protein 4.1R (tan) at the spectrin-actin interaction sites (5). Spectrin tetramers are depicted in the folded conformation (~60nm) of the native, unspread membrane skeleton, but are able to unfold and extend to nearly 200nm in length (5, 7).

## Materials and Methods

Experimental procedures for culturing human CD34+ cells, flow cytometry and qRT-PCR; immunofluorescence labeling of NMIIA and phalloidin labeling of F-actin in RBCs; Airyscan and TIRF microscopy and image analysis; RBC isolation and biochemical procedures; blebbistatin treatment of RBCs; confocal microscopy and 3D shape measurements; membrane flickering and microfluidics assays, are described in *S1 Materials and Methods*.

## Acknowledgements

We thank Connie Smith and Ilan Wittstein for helping V.M.F. with the initial biochemical experiments, and the members of the Fowler lab for helpful suggestions and comments. This work was supported by a grant from the Whitaker Foundation to V.M.F., National Institutes of Health Grants GM34225 and HL083464 (V.M.F.), HL134043 (Subcontract to L.B.), and HL126497 (I.G.), and a National Science Foundation award (CBET – 1560709) to J.W.. A.S. was supported by an NIH/NCATS CTSA Award UL1 TR001114 to the Scripps Translational Science Institute. L.B. is the recipient of an Allied World St. Baldrick’s Scholar.

## Supplemental Figure Legends

**Figure S1.**
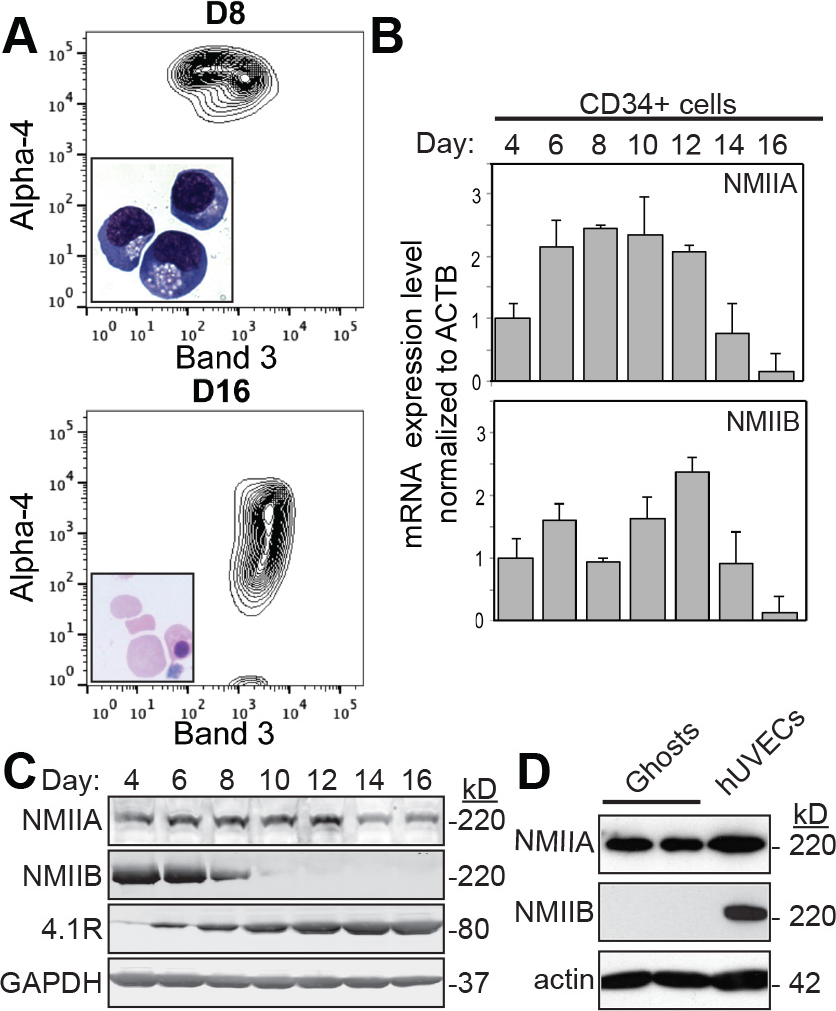
NMIIA HC (*MYH*9) is the predominant RBC NMII isoform expressed during human erythroid terminal differentiation. (A) Flow cytograms at day 8 (upper panel) and day 16 (bottom panel) of *in vitro* erythroid differentiation of CD34^+^ cells demonstrate that α4-integrin levels decrease while Band 3 levels increase over time, as expected for differentiation towards RBCs. Insets show H&E-stained cells, most of which have enucleated by day 16. (B) qRT-PCR of *MYH9* (NMIIA HC) and *MYH10* (NMIIB HC) transcript levels normalized to *ACTB* during erythroid differentiation of CD34^+^ cells show that both transcripts decrease during differentiation. (C) Immunoblots of NMIIA HC and NMIIB HC of the same samples demonstrate that NMIIB protein levels decrease during erythroid differentiation while NMIIA HC protein levels persist. As a control for erythroid differentiation, 4.1R expression levels are increased, as expected. GAPDH is used as a loading control. (D) Immunoblot indicating that NMIIA HC but not NMIIB HC is detected in Mg^++^ ghosts prepared from human RBCs from normal donors. Cultured human umbilical vein endothelial cell (hUVEC) lysate was used as a positive control for the presence of NMIIA and NMIIB HCs. Actin is used as a loading control.

**Figure S2.**
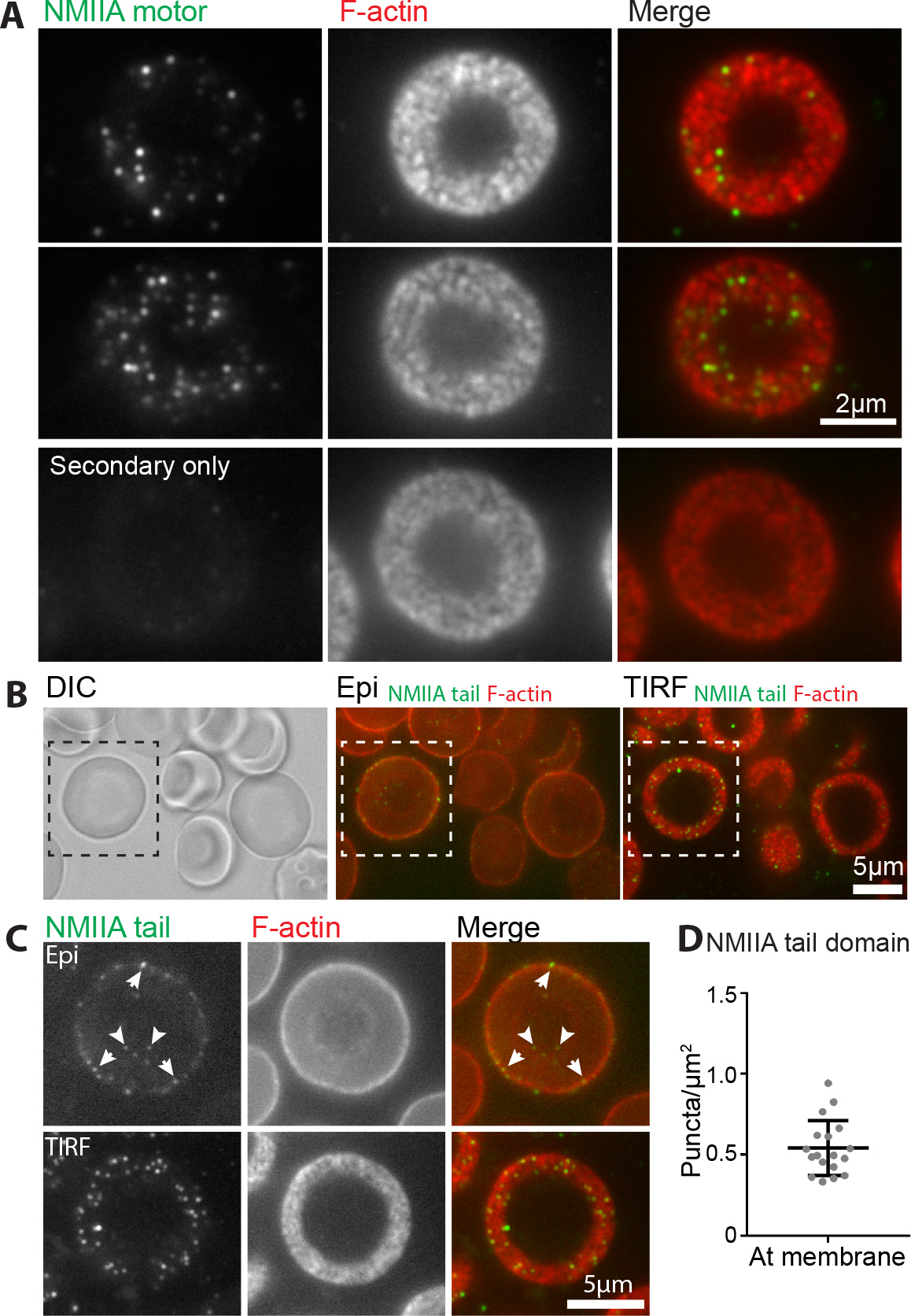
NMIIA tail domains and motor domains show similar distributions at the RBC membrane. (A) Top two rows show TIRF images of RBCs immunostained with an antibody against the NMIIA motor domain (green, as in Figure 1) and rhodamine phalloidin for F-actin (red). Bottom row shows TIRF images of RBCs immunostained with only secondary antibody (green) and rhodamine phalloidin (red). The minimal signal in the green channel demonstrates that the primary antibody-stained NMIIA puncta in RBCs represent specific staining of NMIIA. (B) Differential interference contrast (DIC), epifluorescence, and TIRF images of a group of RBCs immunostained with an antibody against the NMIIA tail domain (green) and rhodamine phalloidin for F-actin (red). (C) Higher-magnification views of the boxed cell in (B). In the epifluorescence image (top), NMIIA puncta are at the membrane at the rim of the cell (arrows) and the dimple at the center of the cell (arrowheads). In the TIRF image (bottom), NMIIA staining is distributed in puncta at the membrane, similar to (A) and Figure 1. (D) Quantification of the density of NMIIA puncta/μm^2^ stained with the antibody against the tail domain within 200nm of the membrane from TIRF images. n = 18 RBCs from 1 individual donor. Lines represent mean ± S.D. (0.54 ± 0.17 puncta/μm^2^, similar to the value in Figure 1G).

**Figure S3.**
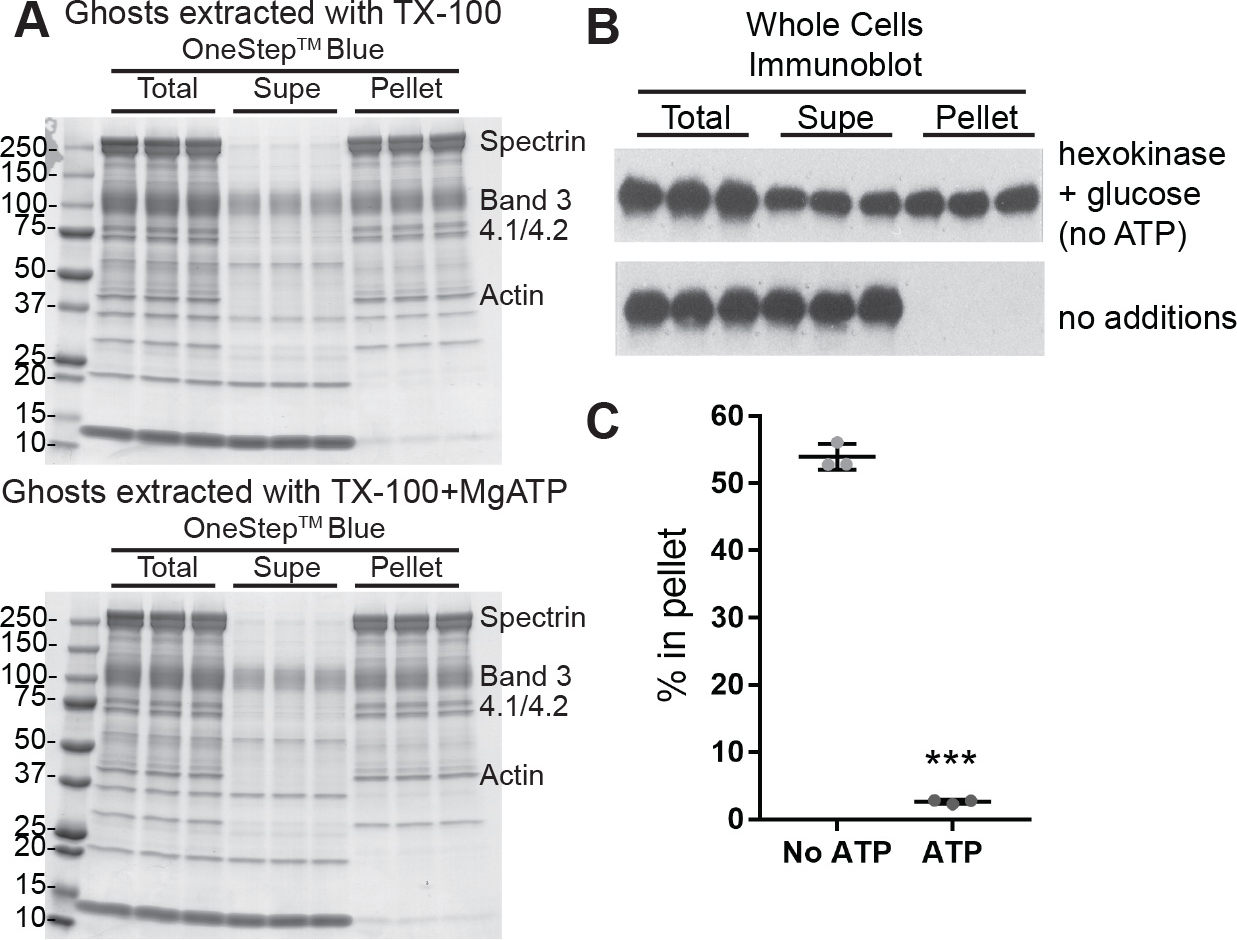
The association of NMIIA with the RBC membrane skeleton is ATP-dependent. (A) SDS-PAGE gels of samples from Triton X-100 extraction of ghosts in Figure 2A-C, stained with OneStep™ Blue. No disassembly of the Triton X-100 insoluble membrane skeleton is observed in the samples extracted with MgATP (bottom) when compared to samples extracted without MgATP (top). (B) Immunoblots of NMIIA HC after Triton X-100 extraction of whole RBCs shows that NMIIA association with the membrane skeleton is MgATP-dependent, in agreement with the results in ghosts shown in Figure 2A-C. RBCs were lysed in TX-100 and depleted of endogenous ATP with hexokinase and glucose before sedimentation and SDS-PAGE on large format 7.5% acrylamide gels, followed by immunoblotting (see Methods). Unlike the small format gels used in Figure 2A, the mobility of the NMIIA HC is identical in allsamples, since these large format gels resolve the NMIIA HC from the spectrin bands (not shown). (C) Quantification shows that RBCs lysed in TX-100 and depleted of ATP retain 53.9 ± 1.9% NMIIA in the Triton X-100-insolubule pellet, while cells with endogenous ATP levels retain only 2.6 ± 0.3% NMIIA in the Triton X-100-insoluble pellet. Each point represents one of three technical replicates from one individual donor. Data reported as mean ± S.D. p < 0.0001.

**Figure S4.**
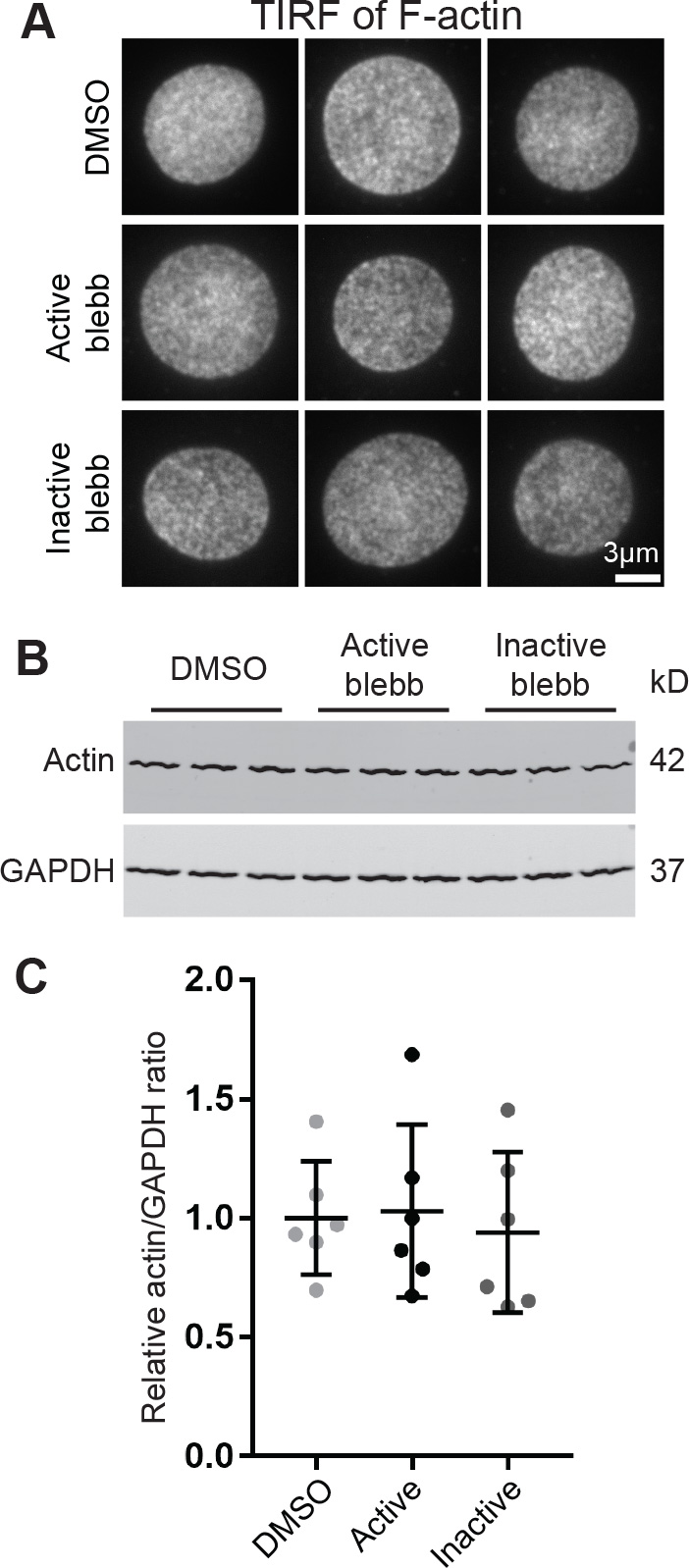
NMIIA motor activity does not control the micron-scale distribution of F-actin at the RBC membrane or the RBC soluble actin pool. (A) Representative TIRF images of RBCs treated with DMSO, 20μM active blebbistatin, or 20μM inactive blebbistatin as in Figure 3. No differences in the dense, reticular distribution of F-actin are detectable at this resolution. (B) Western blots of Triton-X-100 supernatants from RBCs treated with DMSO alone, 20μM active blebbistatin, or 20μM inactive blebbistatin. (C) Quantification of the ratios of actin to GAPDH. Each point represents one of three technical replicates from two individual donors for a total of n = 6. Data reported as mean ± S.D. There is no significant difference between treatment groups.

**Figure S5.**
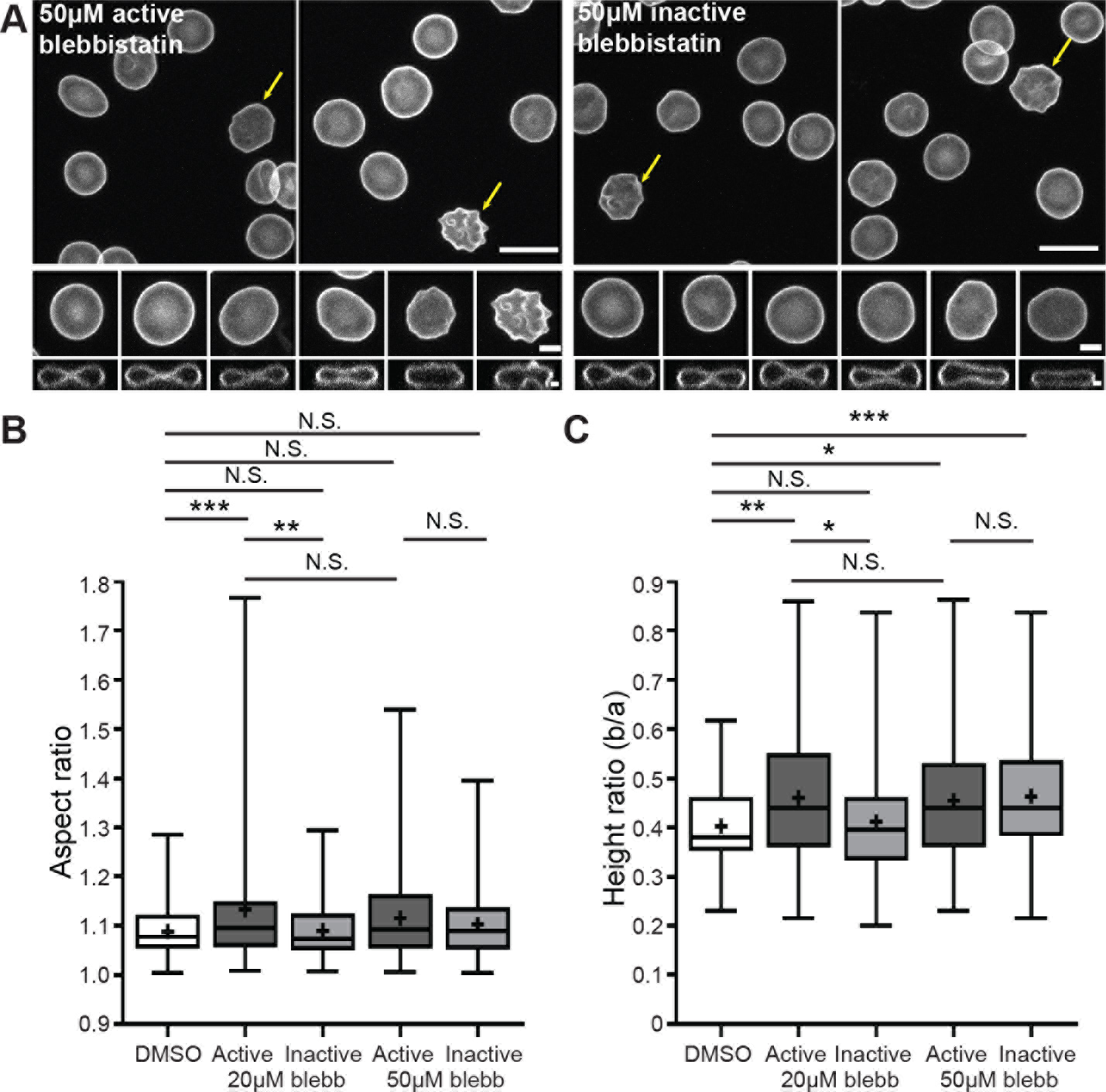
Dose-dependent effects of blebbistatin treatment on RBC morphology. (A) Confocal images of GPA-stained RBCs after treatment with 50μM active or 50μM inactive blebbistatin. Top and middle rows show XY maximum-intensity projections, while the bottom row shows XZ slices from the middle of the cell. Note occasional crenated early-stage echinocytes (arrows) and cells with relaxed dimples after treatment with both 50μM active and 50μM inactive blebbistatin. (B) Box-and-whisker plots showing the distributions of aspect ratios (RBC elongation) for RBCs treated with DMSO, or either 20μM or 50μM active or inactive blebbistatin. In contrast to results with 20μM blebbistatin, RBCs treated with 50μM active and inactive blebbistatin do not have an increased aspect ratio compared to RBCs treated with DMSO. (C) Box-and-whisker plots showing the distributions of height ratios (RBC biconcavity) for RBCs treated with DMSO or either 20μM or 50μM active or inactive blebbistatin. RBCs treated with 50μM active or inactive blebbistatin both have increased height ratios, suggesting that this change is due to off-target effects of blebbistatin at this concentration. By contrast, 20μM active but not 20μM inactive blebbistatin leads to an increased RBC height ratio, indicating that the loss of RBC biconcavity at this concentration is due to specific inhibition of myosin activity. For box and whisker plots, middle line represents median, upper and lower lines represent third and first quartiles respectively, and whiskers represent minimum and maximum values. + sign denotes means. DMSO, n = 107; 20μM active blebbistatin, n = 114; 20μM inactive blebbistatin, n = 104; 50μM active blebbistatin, n = 107; 50μM inactive blebbistatin, n = 102 cells from one donor.

**Figure S6.**
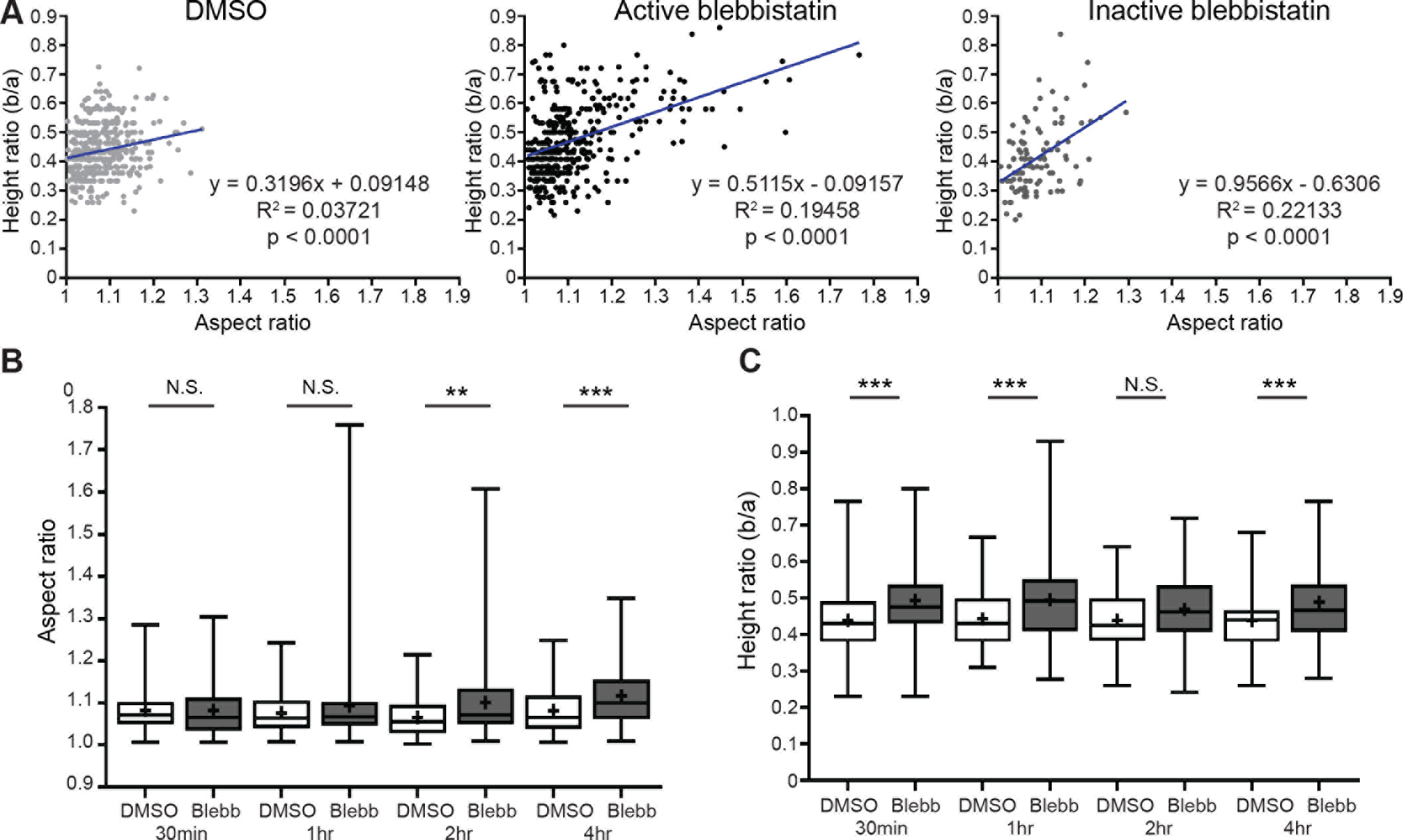
Relationship between RBC elongation (aspect ratio) and biconcavity (height ratio). (A) Scatterplots of aspect ratio versus height ratio for RBCs treated with DMSO, 20μM active blebbistatin, or 20μM inactive blebbistatin. Blue lines represent linear trendlines. The equation for the best-fit line, the R^2^ value, and the p-value for each linear regression are given next to the chart. In all of the treatment groups, height ratio generally increases with increasing aspect ratio, though there is high variability in this trend between individual cells. (B-C) The time courses of RBC elongation and loss of biconcavity due to blebbistatin treatment are asynchronous. (B) Box and whisker plots showing the distributions of aspect ratios for cells treated with DMSO or 20μM active blebbistatin for 30 minutes, 1 hour, 2 hours (p = 0.001), or 4 hours (p = 0.0007). No significant changes were observed for DMSO-treated cells at any time point by one-way ANOVA. (C) Box-and-whisker plots showing the distributions of height ratios for the RBCs in B. 30 minutes p = 0.0002, 1 hr p = 0.0006, 4 hr p = 0.0006. No significant changes were observed for DMSO-treated cells at any time point by one-way ANOVA. While the aspect ratios are not significantly different until 2 hours of treatment with blebbistatin, the height ratios increase by 30 minutes. For box and whisker plots, middle line represents median, upper and lower lines represent third and first quartiles respectively, and whiskers represent minimum and maximum values. + sign denotes means. 30min DMSO n = 98; 30min blebbistatin, n = 104; 1hr DMSO, n = 95; 1hr blebbistatin, n = 103; 2hr DMSO, n = 99; 2hr blebbistatin, n = 100; 4hr DMSO, n = 100; 4hr blebbistatin, n = 109 cells from one individual donor.

**Figure S7.**
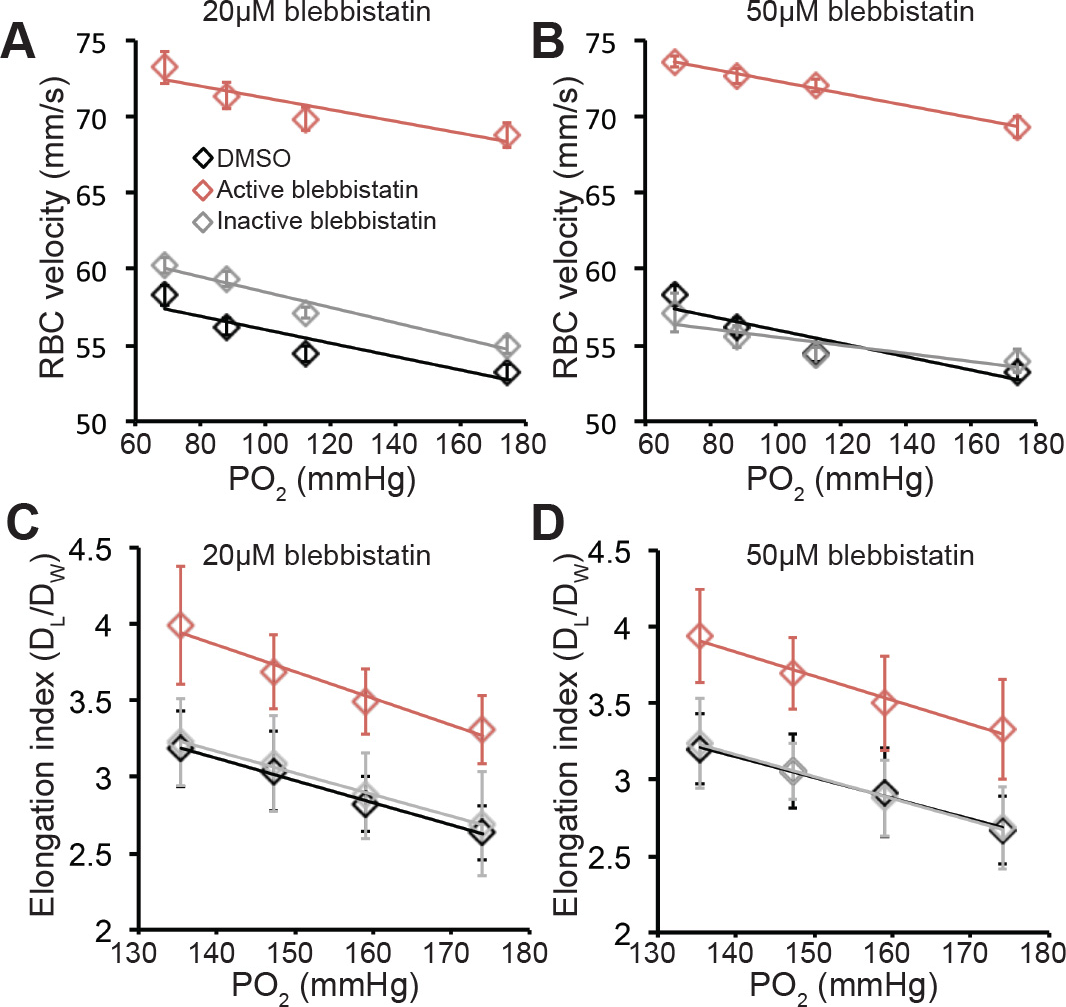
Blebbistatin treatment does not affect the dependence of RBC deformability on decreasing levels of oxygen. (A-B) Relationship between RBC velocity in 5μm-wide channels and oxygen concentration for RBCs treated with DMSO or active or inactive blebbistatin at 20μM (A) or 50μM (B). Data are represented as mean ± S.D. (A) Trendlines: DMSO, y = −0.0441x + 60.421; active blebbistatin, y = −0.0387x + 75.096; inactive blebbistatin, y = −0.0504x + 63.524. (B) Trendlines: DMSO, y = −0.0441x + 60.421; active blebbistatin, y = −0.0403x + 76.374; inactive blebbistatin, y = −0.0267x + 58.205. (C-D) Relationship between RBC elongation index in 20μm-wide constrictions in 30μm-wide channels and oxygen concentration for RBCs treated with DMSO or active or inactive blebbistatin at 20μM (C) or 50μM (D). Data are represented as mean ± S.D. (C) Trendlines: DMSO, y = −0.0145x + 5.157; active blebbistatin, y = −0.0175x + 6.3073; inactive blebbistatin, y = −0.0141x + 5.1461. (D) Trendlines: DMSO, y = −0.0136x + 5.0513; active blebbistatin, y = −0.0157x + 6.038; inactive blebbistatin, y = −0.0144x + 5.1759. In (A-D), each point represents the mean of 30-48 RBCs from two or three individual donors.

## S1 Materials and Methods

### CD34^+^ cell isolation and erythroid culture

CD34^+^ cells from cord blood were isolated by positive selection, using the Magnetic Activated Cell Sorting (MACS) beads system (Miltenyi Biotec, San Diego, CA), according to the manufacturer’s instructions. CD34^+^ cells were then induced to erythroid differentiation using a 3-phase liquid culture system over a period of 18 days as described previously (1). Flow cytometry was used to monitor erythroid differentiation, with Glycophorin A (GPA), Band 3 and α4-integrin as surface markers. Fluorescence intensity was evaluated using a BD Fortessa (Becton Dickinson) flow cytometer, and data were subsequently analyzed using FlowJo v10.3 (Tree Star Inc.).

### qRT-PCR

RNA was isolated every other day from day 4 to day 16 of erythroid differentiation. RNA isolations were performed following manufacturer’s instructions (RNeasy Plus, Qiagen). Briefly, cDNA was generated from 1 μg of RNA using SuperScript VILO Master Mix (Life technologies), and *MYH9* or *MYH10* gene expression assessed by quantitative PCR using a QuantiStudio 3 Real-Time PCR system (Life Technologies). Gene expression levels were analyzed in triplicate, and target mRNA levels were normalized to *ACTB* expression. Relative changes in gene expression were determined by the following equation: Efficiency^ΔCt reference^/Efficiency ^ΔCt target^. Primers used were as follows: *MYH9,* Forward: 5′-CACTGAGACGGCCGATGC-3′ Reverse: 5′-GTCCCCGCGCCTGAG-3′; *MYH10,* Forward: 5′-CCTGCGAACCCTCCTGGT-3′ Reverse: 5′-TTTTATCTGTGGCTTTAGGGAACC-3’; ACTB, Forward: 5’-AGGCACCAGGGCGTGAT-3’

### Isolation of RBCs and preparation of Mg^++^ ghosts

Whole blood was collected from healthy human donors into EDTA tubes (BD Diagnostics). Blood was added to 5 volumes of ice-cold PBS/Dextran (150mM NaCl, 5 mM NaHPO_4_ pH 7.4, 0.75% dextran [Sigma 31392], 0.02% NaN_3_), then settled at 1 × *g* to separate the RBCs from white blood cells and platelets. The RBC pellet was washed four times in ice-cold PBS (150 mM NaCl, 10 mM NaHPO_4_, pH 7.4) by centrifuging for 5 min at 600 × *g* and resuspension in PBS. Packed RBCs were then lysed in 40 volumes of ice-cold hypotonic lysis buffer containing magnesium ( (5 mM NaHPO_4_ pH 7.4, 2 mM MgCl_2_, 1 mM DTT, 20 μg/ml PMSF) while vortexing vigorously, followed by centrifuging for 10 min at 39,000 × *g* at 4°C to sediment the RBC membranes (Mg^++^ghosts) (2). The pellet was resuspended in ice-cold lysis buffer while vortexing, and membranes sedimented as above, for a total of 4 washes, to remove hemoglobin and other cytosolic proteins. Ghosts were then aliquoted, flash-frozen in liquid nitrogen, and stored at −80°C until use.

### Preparation of membrane skeletons by TX-100 extraction of Mg^++^ ghosts

Five volumes of ice-cold TX-100 extraction buffer (0.2% TX-100 (Sigma), 2 mM MgCl_2_, 1 mM EGTA, 10 mM NaHPO_4_ pH 7.5, 1 mM DTT, 1:2000 protease inhibitor cocktail [Sigma P8340], and phosphatase inhibitor cocktail [Thermo Scientific product #88667]) was added to a 50μ aliquot of Mg^++^ ghosts. In some experiments, the TX-100 extraction buffer also contained 5mM MgATP. The sample was then vortexed and incubated on ice for five minutes. A small volume was removed to make the “Total” gel sample. The remainder was centrifuged at 21,000 × *g* at 4°C to sediment the TX-100-insoluble membrane skeletons. A small volume of the supernatant was removed to make the “Supe” gel sample, and the rest of the supernatant was discarded. The TX-100-insoluble pellet was resuspended by vortexing and sonication in sufficient TX-100 extraction buffer to obtain the volume of the original sample after removal of the aliquot for the “Total” gel sample. A small volume of the resuspended pellet was used to make the “Pellet” gel sample. Gel samples were prepared by mixing with equal volumes of 2× SDS sample buffer (210mM Tris-Cl, pH 6.8, 3mM EDTA, 20% sucrose, 0.01% bromophenol blue, 6% SDS, 85mM DTT) and boiling for five minutes.

### SDS-PAGE and Immunoblotting

Gel samples of ghosts and membrane skeletons were run on Novex™ WedgeWell™ 4-20% Tris-glycine gels (Invitrogen) with a 4% stacking gel at 150 V for 75 minutes. The gels were then either stained with One-Step Blue protein gel stain (Biotium) or prepared for immunoblotting. Proteins were transferred onto 0.2 μm nitrocellulose (Genesee Scientific) at 120 V for 1 h at 4°C. For actin and other low-molecular-weight proteins, the transfer buffer contained 20% methanol, 12.5 mM Tris, and 96 mM glycine (3). To transfer the NMII heavy chain, the transfer buffer also contained 0.1% SDS. Transfer buffers were pre-cooled to 4°C, and blue ice packs were placed in the apparatus, and changed if necessary, to prevent overheating. After transfer, nitrocellulose membranes were incubated in DPBS for 1 h at 65°C and then stained with ATX Ponceau S red staining solution (Sigma 09189) to assess the quality of the transfer. Membranes were blocked in 4% BSA in PBS for 2 h at room temperature and then incubated with primary antibodies overnight at 4°C in the cold room. Membranes were washed four times in PBS + 0.1% TX-100, then incubated with secondary antibody at room temperature for 1 h, and then washed four times in PBS + 0.1% TX-100. Primary and secondary antibodies were diluted into 40 mg/ml BSA, 150 mM NaCl, 10 mM NaHPO_4_ pH 7.5, 1 mM EDTA, 0.2% TX-100. Primary antibodies used were: mouse anti-NMIIA tail domain antibody (Abcam ab55456), rabbit anti-pS1943 NMIIA (Cell Signaling Technologies 5026), rabbit anti-GAPDH (Santa Cruz Biotechnology sc-25778), and mouse anti-actin (Chemicon MAB1501). All antibodies were diluted 1:1000 except for the actin antibody, which was diluted 1:10000. Secondary antibodies used were goat anti-mouse IRDye 680LT (LI-COR 926-68020) and goat anti-rabbit IRDye 800CW (LI-COR 926-32211) at a dilution of 1:10000. Labeled bands on blots were imaged on a LI-COR Odyssey Infrared Imaging System, and background-corrected intensity measurements of protein bands were measured using ImageJ.

### Preparation of membrane skeletons by TX-100 extraction of whole RBCs

RBCs were isolated from whole blood by sedimentation through Dextran/PBS as described above, washed three times in 10 mM HEPES, pH 7.4, 150 mM NaCl, 0.1 mM EGTA, and resuspended to 20% vol/vol in the same buffer on ice. Cells were lysed by addition of an equal volume of ice cold (a) 2.0% TX-100, 20 mM HEPES, pH 7.3, 150 mM KCL, 5 mM EGTA, 4 mM MgCl_2_. 1 mM DTT and 40 μg/ml PMSF, or (b) the same buffer containing 25 U/ml hexokinase (Sigma H-4502) and 2 mM glucose to deplete endogenous ATP. Cell suspensions were incubated 30 min at 37°C, membrane skeletons were collected by centrifugation for 20 min at top speed in an Eppendorf microfuge at 4°C, the supernatants were removed, and pellets resuspended to the original volume in the lysis buffer. SDS gel samples were prepared of the total lysate before centrifugation, the supernatant, and membrane skeleton pellets. Equivalent amounts were subjected to SDS-PAGE on large format 5% acrylamide gels (Hoefer), and transferred to nitrocellulose for 3 h at 15°C in pre-cooled 20% methanol, 12.5 mM Tris, and 96 mM glycine plus 0.1% SDS. NMIIA heavy chain bands were detected by labeling with 10 μg/ml affinity-purified rabbit antibodies to human platelet myosin (4) as described above, except the primary antibody was detected with ^125^I-Protein A (2 × 10^6^ cpm/ml) as described (2). The percent of NMIIA in the total, supernatant and pellet fractions was determined by excising the ^125^I-Protein A-labeled bands from the nitrocellulose and counting in a Beckman gamma counter. Nonspecific labeling was corrected by subtracting counts from a similar sized region of nitrocellulose not containing NMIIA.

### ^32^P-orthophosphate metabolic labeling of RBCs and immunoprecipitation (IP) of NMIIA

RBCs were isolated from whole blood by sedimentation through Dextran/PBS as described above, washed three times in 10 mM HEPES, pH 7.4, 150 mM NaCl, followed by three times in 25 mM NaHCO_3_, pH 7.4, 120 mM NaCl, 3.7 mM KCl, 2.4 mM MgCl_2_, 1.2 mM CaCl_2_, and then resuspended to 20% vol/vol in the same buffer containing 1 mM adenosine (Sigma), 1 mM inosine (Sigma) and 10 mM glucose (Baker), 100 mg/ml penicillin G and streptomycin (Sigma). 80 μl of 10 mCi/ml ^32^P-orthophosphoric acid (1 mCi/ml packed RBCs) (New England Nuclear, Boston MA) was added to the cell suspension and incubated with shaking for 24 h at 37°C. The ^32^P-labeled RBCs were washed four times in 10 mM Tris, pH 7.5, 150 mM NaCl, 0.02% Na_3_, resuspended to 20% vol/vol in the same buffer and lysed in an equal volume of 2× Triton-Lysis IP buffer modified from Berlot et al (5) (2% TX-100, 0.2 M NaF, 40 mM sodium pyrophosphate, 10 mM EDTA, 0.2 M KH_2_PO_4_ and 40 mM Tris, pH 7.5, 0.02% NaN_3_) containing 10 μg/ml PMSF and 0.5 mM iodoacetic acid, and then incubated 20 min on ice. The RBC lysate was centrifuged for 20 min at 18,000 rpm in a JA20 rotor at 2°C to remove the insoluble membrane skeleton. SDS-PAGE samples were prepared of the whole RBC lysate and the lysate supernatant after centrifugation (IP input). To IP myosin, 1 ml of each supernatant sample (derived from a 10% vol/vol RBC suspension) was added to microfuge tubes containing Staphylococcus A bacteria (Staph A; Pansorbin, Calbiochem), to which antibodies to human platelet myosin (2.6 μg affinity-purified rabbit IgG; (4) or pre-immune rabbit IgG (2.6 μg) had been pre-adsorbed for 2 h at 4°C in 1× Triton-Lysis IP buffer containing 1 mg/ml BSA. Samples were incubated overnight at 4°C on an end-over-end rotator, and unbound proteins were removed by washing the Staph A three times in 1× Triton-Lysis IP buffer plus 1 mg/ml BSA, two times in Triton-Lysis IP buffer, and two times in 50 mM Tris, pH 7.5 using a microfuge at 4°C. Immunoprecipitated proteins were solubilized from the Staph A in 1× SDS sample buffer by boiling for 2 min, centrifuged briefly to sediment the Staph A, and then samples were electrophoresed on a 5-15% linear acrylamide gradient SDS gel containing 4 M urea with a 5% stacking gel containing 2 M urea (2). The gel was stained with Coomassie brilliant blue R-250, dried onto Whatman filter paper using a gel vacuum dryer, and then exposed to Kodak XAR-5 x-ray film at −80°C with an intensifying screen (Dupont Lightning Plus) to detect ^32^P-labeled bands.

### Blebbistatin treatment of RBCs

Whole blood was collected as above and RBCs were washed three times using hHBS (145 mM NaCl, 5 mM KCl, 1 mM MgCl_2_, 10 mM glucose, 10 mM HEPES, pH 7.4, with 2 mM adenosine added on the day of the experiment). Packed RBCs were resuspended in hHBS (1% v/v). DMSO, inactive R-blebbistatin (Cayman Chemical 13165), or active S-blebbistatin (Cayman Chemical 13013) was added to the RBC suspension to a final DMSO concentration of 0.1%. RBCs were incubated at 37°C for 2 h in a shaking water bath adjusted to keep the RBCs in suspension and well-mixed during the incubation.

To measure the relative levels of actin in the cytosol following blebbistatin treatment, RBCs were lysed by vortexing in four volumes of TX-100 lysis buffer (145mM NaCl, 5mM KCl, 1mM MgCl_2_, 10mM HEPES pH 7.4, 2.5% TritonX-100, 2mM EGTA, 5mM DTT, 1:2000 protease inhibitor cocktail (Sigma P8340)). The RBC lysate was underlaid with 10% sucrose in the TX-100 lysis buffer. The lysate was then centrifuged at 21,000 × *g* at 4°C to sediment the TX-100-insoluble membrane skeletons. Gel samples were prepared by mixing a small volume of the supernatant with ¼ volume of 5× SDS sample buffer (540mM Tris-Cl, pH 6.8, 7mM EDTA, 50% sucrose, 0.04% bromophenol blue, 11% SDS, 700mM DTT) and boiling for five minutes.

### Immunofluorescence staining of RBCs

Whole blood was collected from healthy human donors into EDTA tubes as above (BD Diagnostics). For NMIIA immunostaining and rhodamine-phalloidin staining, 20 μl of whole blood was added to 1 ml of 4% paraformaldehyde (PFA, Electron Microscopy Sciences) in Dulbecco’s PBS (DPBS – Gibco) mixed, and incubated at room temperature overnight. Fixed RBCs were washed three times in DPBS by centrifuging for 5 minutes at 1000 × *g*, permeablized in DPBS + 0.3% TX-100 for 10 minutes, and then blocked in 4% BSA, 1% normal goat serum in DPBS (Blocking Buffer, BB) at 4°C for at least 4 days or up to 1 week before immunostaining. Permeabilized and blocked RBCs were then incubated with rabbit anti-NMIIA motor domain antibody (Abcam ab75590) or mouse anti-NMIIA tail domain antibody (Abcam ab55456) diluted in BB for 2-3 h at room temperature, washed two times in BB as above, and then incubated in Alexa-488-conjugated goat anti-rabbit secondary antibody (Life Technologies A11008) or goat anti-mouse secondary antibody (Life Technologies A11001) mixed with rhodamine-phalloidin (Life Technologies R415 at a final concentration of 130nM) in BB for 1-2 h at room temperature, followed by washing three times in BB as above. Stained cells were cytospun onto coverlips for TIRF microscopy or slides for confocal microscopy and mounted with ProLong™ Gold mounting medium (Invitrogen) prior to imaging.

For TIRF imaging of RBCs after treatment with blebbistatin, ~1μl packed RBCs were resuspended in 4% PFA in DPBS and incubated at room temperature for four hours. Fixed RBCs were washed three times in DPBS by centrifuging for 5 minutes at 1000 × *g*, permiablized in DPBS + 0.3% TX-100 for 10 minutes, then blocked in 4% BSA, 1% normal goat serum in BB overnight. Permeabilized and blocked RBCs were then incubated with rabbit anti-NMIIA motor domain antibody (Abcam ab75590) diluted in BB for 2-3 h at room temperature, washed two times in BB as above, and then incubated in Alexa-488-conjugated goat anti-rabbit secondary antibody (Life Technologies A11008) mixed with rhodamine-phalloidin (Life Technologies R415 at a final concentration of 130nM) in BB for 1-2 h at room temperature, followed by washing three times in BB as above. Stained cells were cytospun onto coverlips for TIRF microscopy.

For RBC shape measurements after blebbistatin treatment (Figure 4), the RBC suspension was mixed with an equal volume of 8% PFA (final PFA concentration of 4%), and fixed overnight at room temperature. RBCs were then washed three times in DPBS by centrifugation as above, blocked for 1 h in 4% BSA, 1% normal goat serum in DPBS, and stained with the FITC-conjugated mouse anti-GPA antibody (BD Pharmingen 559943) for 1 h at room temperature. GPA-stained RBCs were cytospun onto glass slides and mounted with ProLong™ Gold mounting media and coverslipped prior to imaging.

### Fluorescence Microscopy

RBCs immunostained for NMIIA and rhodamine phalloidin for F-actin were imaged using a Nikon Eclipse Ti inverted microscope with a 100× Apochromat oil objective (NA 1.49), either by epifluorescence microcsopy (Figure S2B) using vertical illumination with 488 and 561 laser lines and an ORCA-Flash 4.0 V2 Digital CMOS camera (Hamamatsu) or by TIRF illumination with the same laser lines (Figure 1F). Images were acquired using NIS-Elements 4.1 software. RBCs immunostained for NMIIA and rhodamine phalloidin for F-actin were also imaged using a sensitive Zeiss LSM 880 Airyscan laser scanning confocal microscope with a 63× or 100× 1.46 NA oil Plan Apo objective (Figure 1A-C). Z-stacks were acquired at a digital zoom of 1.8 and a Z-step size of 0.168 μm. Note that standard confocal laser scanning microscopy was unable to image NMIIA immunostaining in RBCs due to the dim signal, relatively low sensitivity of standard confocal microscopy, and photobleaching during confocal image acquisition.

RBC shape measurements of GPA-stained RBCs (Figure 4) were obtained from confocal Z-stacks acquired on a Zeiss LSM 780 laser scanning confocal microscope using a 100× oil Plan Apo objective with 1.4 NA. Z-stacks were acquired at a digital zoom of 1.0 and a Z-step size of 0.25 μm. RBC aspect ratios were measured from maximum intensity projections of Z-stacks using ImageJ. Height measurements were acquired manually from XZ views of the center of each RBC in Volocity.

### Image analysis

Numbers of NMIIA puncta at the membrane in immunostained RBCs (Figure 1F-G and Figure 3) were counted automatically from TIRF images in Volocity (Quorum Technologies) using the “Find Spots” function in the “Measurements” module. The number of NMIIA puncta per RBC and the minimum distances between puncta (Figure 1C-E) were determined from 3D reconstructions of AiryScan Z-stacks in Imaris (BitPlane) using the “Spot to Spot Minimum Distance” function in the “Spot Identification” module. Puncta were only counted if they were within the area of rhodamine-phalloidin (F-actin) staining, above the maximum intensity observed in RBCs stained with secondary antibody alone, and imaged at the same settings. To correct for Z-stretch, the distance between Z-steps was set to 0.085 μm. For RBC shape measurements (Figure 4), aspect ratios were measured from maximum-intensity projections of Z-stacks using ImageJ. Height measurements were acquired manually from XZ views of the center of each RBC in Volocity. To correct for Z-stretch, the distance between Z-steps was set to 0.18 μm prior to recording height measurements.

### Flickering measurements

Spontaneous RBC membrane oscillations (flickering) were measured as previously described (6). Positive-low, phase-contrast, time-lapse images of RBCs seeded on microscope slides were recorded for 10 s at a rate of 33 frames/s using a 100× UPlanApo phase contrast objective (NA 1.35) on an Olympus BX62 microscope. Flickering was measured using iVision 4.0.9 software (BioVision Technologies). At the end of each recording, an intensity projection step of the image stack was performed to identify and exclude RBCs that drifted during recording. The intensity of scattered light was used to calculate, pixel by pixel, the coefficient of variance at each point within the RBC and display the results as a pseudocolor amplitude map.

### Microfluidics assays

Polydimethylsiloxane (PDMS) (Dow Corning Sylgard 184 Silicone Elastomer) microfluidic chips were fabricated using standard single-layer photolithography techniques and experiments performed as described (7). The channel inside the PDMS chip has an overall height of 6.38 μm, and a width of 5 μm at constriction section, mimicking the diameter of capillaries. The microfluidic channel was submerged in the reservoir with various concentrations of sodium sulfite (Sigma-Aldrich) solution (0 M, 0.01 M, 0.1 M and 1 M) in the sink. Because PDMS is permeable to oxygen, the oxygen level inside the microfluidic channel could be tuned from 68 mm Hg to 174 mm Hg (calibrated colorimetrically using Tris(2,2’-bipyridyl)dichlororuthenium(II) hexahydrate (RTDP; Sigma-Aldrich)). A high-speed camera (Phantom) was mounted onto an inverted microscope (63× oil-immersion objective, Leica Microsystems, DMI 6000) to visualize the cell behavior inside the channel. With a sample rate of 1900 fps, the time interval between each frame was 526.31 μs. By measuring the distance the cell moved between frames, the velocities of RBCs flowing through the constriction were calculated.

To investigate changes in RBC deformability under normoxic or hypoxic conditions, a microfluidic channel with a wider constriction was used. The length and width of the constriction channel were 100 μm and 20 μm, respectively. The overall height of the channel was 30 μm. When RBCs were flowing through the constriction channel, RBCs were stretched due to increased shear stress. The length (*D*_*l*_) and the width (*D*_*w*_) of the RBCs was used to calculate the elongation index (*D*_*l*_/*D*_*w*_). For both experiments, the RBC suspension was driven through the channel by using N_2_ with an input pressure of 1.6 psi.

### Statistical analysis

Data are presented in dot plots as mean ± standard deviation (SD). Differences between means were detected using unpaired t-tests. When more than one comparison was made, means were compared using one-way ANOVAs followed by Tukey’s multiple comparisons test. Statistical significance was defined as p < 0.05. Statistical analysis was performed using GraphPad Prism 7.03 software. Box and whisker plots show median values (horizontal center line), third and first quartiles (top and bottom of boxes), and the minimum and maximum of the data (whiskers), as well as the mean (+ sign).

